# Astrocytes control oxytocin-based maternal behavior via connexin 30

**DOI:** 10.64898/2026.01.09.698569

**Authors:** Isabelle Arnoux, Maïna Garnero, Julien Moulard, Astrid Rollenhagen, Carl Meinung, Nadia De Mota, Peggy Barbe, Pascal Ezan, Anthony Laugeray, Vidian de Concini, Alexis Bemelmans, Inga Neumann, Arnaud Menuet, Catherine Llorens-Cortes, Joachim HR Lübke, Grégory Ghézali, Nathalie Rouach

## Abstract

Prosocial and affiliative behaviors rely on complex neuroendocrine neuronal circuits. Dynamic neuroglial interactions play prominent roles in shaping synaptic networks, whose alterations contribute to psychiatric disorders associated with social deficits. However, a role for astrocytes in regulating maternal behavior remains elusive. Here we show that female mice exhibiting increased oxytocin levels in the hypothalamic supraoptic nucleus upon social interactions with pups display neuroglial structural plasticity and downregulation of astroglial connexin 30, a protein involved in morphological remodeling. We found that impairing astroglial connexin 30 expression alters the structural properties of astrocytes by decreasing their volume and coverage of oxytocinergic synapses in the supraoptic nucleus of female mice. This functionally sets levels of plasma oxytocin and maternal behavior, exacerbating maternal care in virgin pup-naïve female mice deficient for astroglial Cx30. Hypothalamic astrocytes, via connexin 30, are thus key determinants of oxytocin-based maternal behavior, whose alteration is known to impair social skills development in offspring.

## INTRODUCTION

Intense morphological neuroglial remodelling occurs during physiological situations of high hormonal demand in the hypothalamo-neurohypophysial system, in which the supraoptic nucleus (SON) is a major component^1^. Magnocellular neurons of the SON synthesize and release neuropeptide hormones, such as oxytocin (OT) or arginine-vasopressin. Previous work established ultrastructurally that during lactation, parturition and osmoregulation, astrocytic processes specifically retract from oxytocinergic, but not from vasopressinergic magnocellular_neurons^2^. Rapid and reversible process retraction alters synaptic strength at local excitatory synaptic inputs notably through reduced astroglial glutamate clearance^3^. These changes in glutamate levels in turn govern the rhythmic drive of oxytocinergic neurosecretory cells^4^, controlling the release of OT in the brain and the blood^5, 6^. The majority of axon terminals originating from OT-secreting magnocellular neurons project to the posterior lobe of the pituitary gland, thereby reaching the systemic circulation. However, local somatodendritic release of OT and long-range axonal projections targeting various forebrain circuitries have also recently been described^7^. Thus, OT not only controls various fundamental reproductive and osmoregulatory functions, but has also a plethora of modulatory effects on cognition and behavior, including especially sociability and maternal care^8^.

Astrocytes have recently been shown to regulate behavior in health and disease^9, 10^. In particular, astrocytes modulate brain circuits controlling behavioral responses, involved in feeding^11, 12^, sleep^13^ or psychiatric disorders^14^. For instance, calcium-dependent ATP release from astrocytes or astroglial Kir4.1 channels can modulate depressive^15, 16^ or anxiety-like^17^ behaviors in mice via regulation of neurotransmission. However, there is still little evidence supporting a role for astrocytes in the neuroendocrine control of social behaviors. Interestingly, repeated exposure of pup-naïve female rat to pups was shown to induce plastic ultrastructural rearrangement of the SON^18^. Moreover, pup exposure drives maternal responsiveness in pup-naïve female mice via OT signaling in the medial preoptic area (MPOA)^19^, a key hypothalamic node controlling the expression of parental behaviors^20^.

Similarly, dams precociously separated from pups at birth, or those whose interactions with young are prolonged by providing new litters, show respectively halted and extended astrocytic processes withdrawal specifically from SON OT-secreting neurons^21^. Accordingly, pup-deprivation impairs proper maternal behavior and associated SON OT neuronal activity in dams^22^. However, it is still unclear whether such hypothalamic astroglial morphological remodelling is merely an incidental consequence of affiliative interactions or if, on the contrary, it actively contributes to OT-based social behaviors. Furthermore, the molecular mechanisms underlying social-cue induced astrocytic structural plasticity remain elusive.

A key property of astrocytes is to express high levels of gap junction (GJ) proteins, the connexins. Interestingly, we have previously shown that the astroglial GJ subunit connexin 30 (Cx30), the expression of which is activity-dependent^23^ and developmentally regulated^24^, controls postnatal structural maturation of hippocampal astrocytes, particularly the extension, ramification and polarity of astroglial processes^23, 25^. These morphological regulations are independent of GJ-mediated biochemical coupling and have functional impact at the synaptic and cognitive levels. In fact, we showed that astroglial Cx30 tunes hippocampal excitatory synaptic transmission, plasticity and memory^25^. Strikingly, the regulation of Cx30 expression also has developmental consequences, since we recently found that the developmental increase in Cx30 expression controls the closure of the critical period for experience-dependent plasticity in the visual cortex^26^. We here show that hypothalamic astrocytes, through Cx30, are key regulators of OT-based maternal behavior.

## RESULTS

### Hypothalamic SON astrocytes display structural and molecular changes in a model of social experience-dependent structural plasticity

Pup exposure drives plastic reorganization of the rat SON ultrastructure^18^ and facilitates maternal behavioral responsiveness of pup-naïve female mice via activation of the oxytocinergic system^19^. To investigate the effects of social experience-dependent structural plasticity in SON astrocytes, we used a pup sensitization model, in which a virgin pup-naïve female adult mouse was exposed to a pup for 30 minutes. We first tested whether interaction with pups increases OT levels in the SON of virgin pup-naïve female mice (Fig. 1a-d). Using in vivo SON microdialysis, a minimally-invasive technique allowing monitoring of dynamic changes in OT concentrations in brain extracellular fluid, we indeed found enhanced OT levels upon pup exposure (n = 14, p = 0.0128, Friedman test followed by Dunn’s post hoc test, Fig. 1c, d). We then investigated whether the functional changes in extracellular OT levels were associated with structural alterations in SON astrocytes. We found that social interactions with pups affect the size and the complexity of GFP-labeled astrocytes from GFAP-eGFP mice (Fig. 1e-i). Specifically, both the total process length and the domain volume were decreased (Sum of process length: control: n = 20 astrocytes from 4 mice, pup exposure: n = 28 astrocytes from 4 mice, p = 0.0010, unpaired t-test; Domain: control: n = 19 astrocytes from 4 mice, pup exposure: n = 27 astrocytes from 4 mice, p = 0.0049, Mann Whitney test, Fig. 1f, g), as assessed with 3D reconstruction of astrocytes. Additionally, the complexity of astrocytes, as assessed by branch level and number of intersections, was also reduced (control: n = 21 astrocytes from 4 mice, pup exposure: n = 28 astrocytes from 4 mice, p = 0.0062, Mann Whitney test and p = 0.0084, two-way ANOVA, respectively, Fig. 1h, i). We next explored whether these morphological changes of astrocyte translated into alterations in the coverage of oxytocinergic neurons. Using immunohistochemistry, we measured the coverage of OT-expressing neurons by GFAP-positive astroglial processes in the somatic zone of the SON (Fig. 1j, k). We observed a reduction in GFAP coverage of OT neuron somas, suggesting a retraction of astrocytic processes following pup exposure (control: n = 23 astrocytes from 5 mice, pup exposure: n = 17 astrocytes from 4 mice, p < 0.0001, unpaired t-test, Fig. 1k). We then investigated the expression of key structural determinants of astroglial morphology and coverage of neurons, i.e., glial fibrillary acidic protein GFAP and Cx30^23, 25^. We found that their expression levels in the SON of virgin pup-naïve female mice were also significantly reduced, as shown by quantitative immunohistochemistry (Cx30: control: n = 10 ROI from 3 mice, pup exposure: n = 13 ROI from 4 mice, p = 0.0001, unpaired t-test; GFAP: control: n = 10 ROI from 3 mice, pup exposure: n = 13 ROI from 4 mice, p = 0.0035, unpaired t-test, Fig. 1l, m). Consistent with these data, we found a decreased expression of putative functional Cx30 proteins, as assessed by western blot analysis of membrane Cx30 proteins labeled by surface biotinylation (Cx30: control n = 8 samples, pup exposure n = 5 samples, p = 0.0095 unpaired t test, Supplementary Fig. 1a, b). Similarly, immunoblotting show decreased levels of GFAP (GFAP control n = 8 samples, pup exposure n = 5 samples, p = 0.036 unpaired t test, Supplementary Fig. 1c, d).

**Fig. 1.**
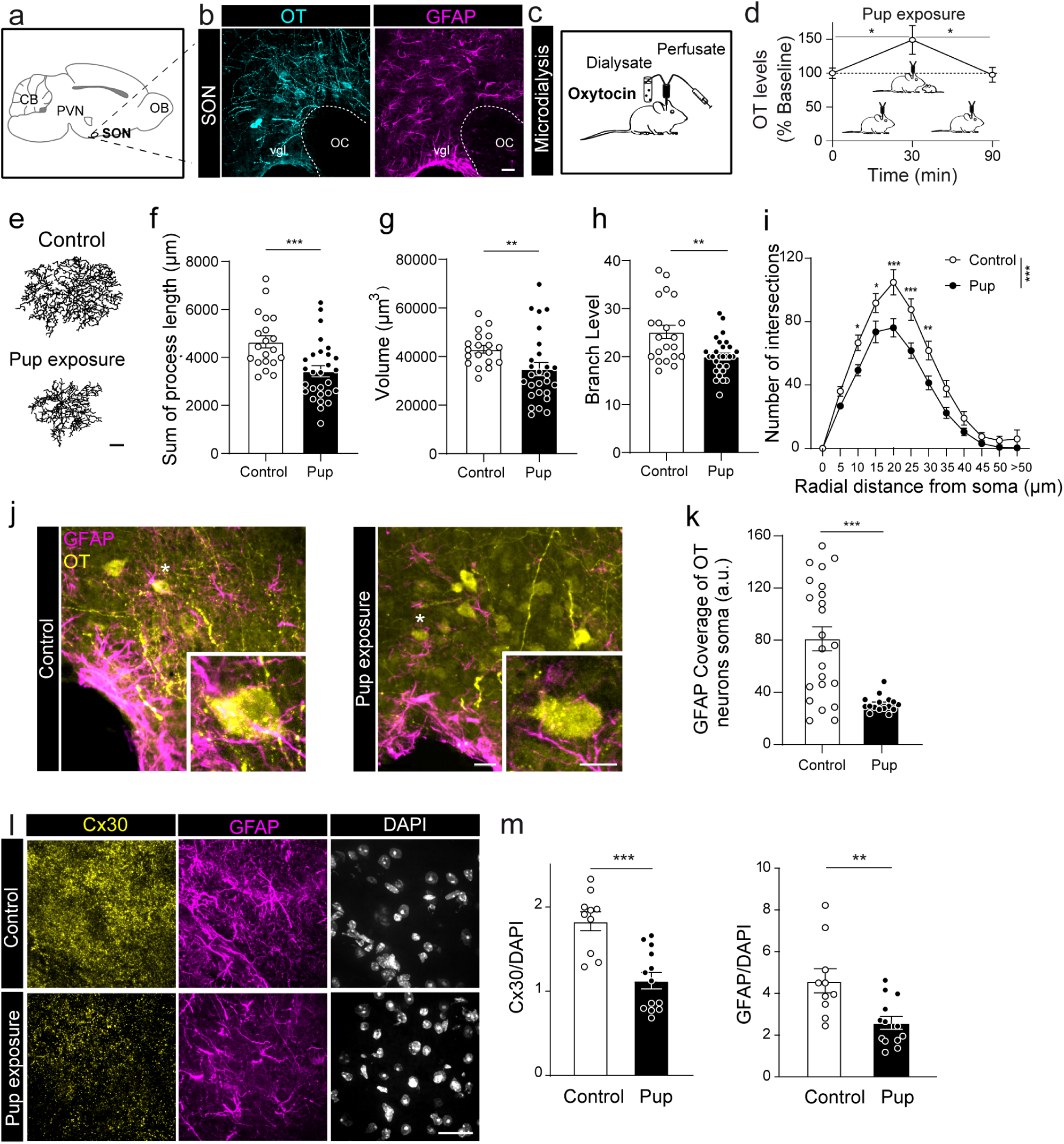
Pup exposure induces OT release and downregulation of Cx30 in the SON of virgin pup-naïve female mice. **a** Scheme depicting localization of the SON. **b** Representative images of OT (cyan) and GFAP (magenta) double-immunofluorescence labeling in the SON of +/+ mice. CB: cerebellum, PVN: paraventricular nucleus, SON: supraoptic nucleus, OB: olfactory bulb, vgl: ventral glia limitans, OC: optic chiasma. Scale bar: 25 µm. **c** Diagram illustrating the in vivo microdialysis method used to measure OT levels in the SON of awake, freely behaving mice. **d** Quantitative analysis of OT levels assessed with microdialysis in +/+ virgin pup-naïve females showing enhanced levels during pup exposure (n = 14, p = 0.0128, Friedman test followed by Dunn’s post hoc test). **e** Three-dimensional reconstruction of astrocytes in the SON of virgin pup-naïve (up) vs pup-sensitized (bottom) female +/+ mice. Scale bar = 10 µm. **f** Sum of process length of astrocytes in the SON of virgin pup-naïve (white, n = 20 astrocytes from 4 mice) vs pup-sensitized (black, n = 28 astrocytes from 4 mice) female +/+ mice (p = 0.0010, unpaired t-test). **g** Domain volume of astrocytes in the SON of virgin pup-naïve (white, n= 19 astrocytes from 4 mice) vs pup-sensitized (black, n = 27 astrocytes from 4 mice) female +/+ mice (p = 0.0049, Mann Whitney test). **h** Branch level of astrocytes in the SON of virgin pup-naïve (white, n = 21 astrocytes from 4 mice) vs pup-sensitized (black, n = 28 astrocytes from 4 mice) female +/+ mice (p = 0.0062, Mann Whitney test). **i** Number of intersections per radial distance from the soma determined by Sholl analysis of astrocytes in the SON of virgin pup-naïve (white, n = 21 astrocytes from 4 mice) vs pup-sensitized (black, n = 28 astrocytes from 4 mice) female +/+ mice (p = 0.0084, two-way ANOVA). **j** Representative images of oxytocinergic neurons (yellow) covered by GFAP-rich astrocyte processes (magenta) in the SON of virgin pup-naïve vs pup-sensitized female +/+ mice. The white asterisks indicate neurons that are magnified in the insets. Scale bar: 20 µm; inset: 8 µm. **k** Pup-sensitized female +/+ mice show reduced GFAP coverage of OT-expressing neurons in the SON (black, n = 17 astrocytes from 4 mice) compared to virgin pup-naïve (white, n = 23 astrocytes from 5 mice, p < 0.0001, unpaired t-test). **l** Representative images showing Cx30 (yellow), GFAP (magenta) and DAPI (white) fluorescent immunostainings in the SON of virgin pup-naïve vs pup-sensitized female +/+ mice. Scale bar: 30 µm. **m** Pup-sensitized female +/+ mice show reduced Cx30 (n = 13 ROI from 4 mice, p = 0.0001, unpaired t-test) and GFAP (n = 13 ROI from 4 mice, p = 0.0035, unpaired t-test) fluorescence intensities compared to virgin pup-naïve controls (n = 10 ROI from 3 mice for both). Fluorescence intensities of Cx30 and GFAP are normalized to DAPI fluorescence intensity. Asterisks indicate statistical significance (*p < 0.05; **p < 0.01; ***p < 0.001).

The decrease in the expression of Cx30 and GFAP was transient, with a significant increase detected one hour after the end of the pup exposure, coinciding with the return of central OT levels to baseline, indicating a dynamic adaptation to the local environment (Cx30: pup 30 min: n = 13 ROI from 5 mice, pup 90 min: n = 27 ROI from 6 mice p < 0.0001, Mann Whitney test; GFAP: pup 30 min : n = 13 ROI from 5 mice, pup 90 min: n = 24 ROI from 6 mice, p < 0.0001, Mann Whitney test, Supplementary Fig 2a-d). The structural changes were also transient, as almost no changes were observed 1h after the end of the pup exposure, except for an increase at the branch level (Sum of process length: pup 30 min: n = 28 astrocytes from 4 mice, pup 90 min: n = 26 astrocytes from 6 mice, p = 0.0003, unpaired t-test; Domain: pup 30 min: n = 27 astrocytes from 4 mice, pup 90 min: n = 25 astrocytes from 6 mice, p = 0.0246, Mann Whitney test; Branch level: pup 30 min: n = 28 astrocytes from 4 mice, pup 90 min: n = 26 astrocytes from 6 mice, p < 0.0001, unpaired t-test; Sholl analysis: pup 30 min : n = 24 astrocytes from 4 mice, pup 90 min: n = 26 astrocytes from 6 mice, p = 0.3767, two-way ANOVA, Supplementary Fig. 2e-i). However, the astrocytic coverage of OT neuron somas remained decreased at this late time point (pup 30 min: n = 15 astrocytes from 3 mice, pup 90 min: n = 12 astrocytes from 3 mice, p = 0.1925, unpaired t-test, Supplementary Fig 2j-k). These data suggest that the changes triggered by pup exposure are reversible and followed a sequence: changes in the expression of Cx30 and GFAP drive the structural remodeling, which ultimately influence the coverage of OT neurons. In the oxytocinergic system, these changes appeared to be specific to the SON, as they were not observed in the paraventricular nucleus (PVN), another important hypothalamic nucleus contributing to OT neurosecretion (Cx30: p = 0.38, unpaired t-test; GFAP: p = 0.1414, unpaired t-test, control: n = 8 ROI from 3 mice, pup exposure: n = 14 ROI from 6 mice; Supplementary Fig. 3). These data suggest that decreased Cx30 expression in the SON of female mice induces structural changes in astrocytes and neuronal coverage.

### Cx30 deficiency reduces astroglial coverage of oxytocinergic neurons and synapses in the SON of virgin females

To test this hypothesis, we took advantage of Cx30 deficient mice (-/-), a mouse model already displaying disrupted Cx30 expression, in which we analysed astroglial coverage of OT neurons in the SON of virgin pup-naïve females. We first measured the extent of OT-expressing neuron coverage by GFAP-positive astroglial processes in the somatic zone of the SON using immunohistochemistry. We found that -/- mice displayed a severe reduction in the coverage of OT neurons by GFAP-positive astroglial processes compared to control mice (perisomatic GFAP intensity (a.u.): +/+: 44.10 ± 2.19, n = 134, -/-: 16.68 ± 1.21, n = 182, p < 0.0001, Mann-Whitney test, Fig. 2a-c). Furthermore, since GFAP occupies only ∼ 15% of the total volume of the astrocyte^27^, we then investigated whether these gross morphological differences reflect changes in OT-positive synapses coverage by fine perisynaptic astroglial processes. To do so, we used high-end fine-scale transmission electron microscopy, and performed quantitative analysis on 3D volume reconstructions of astrocyte-synapse complexes. We observed that the dendritic zone of the SON was composed of dendrites of different calibers occupying most of the neuropil (Fig. 2d, e). The remaining neuropil contained synaptic boutons of variable size, which were established on both, dendritic shafts (Fig. 2e) or spines (Fig. 2d), and were densely packed with synaptic vesicles distributed throughout the terminal (Fig. 2d, e). No qualitative morphological differences of synaptic boutons or dendrites between +/+ and -/- mice could be observed. Measurements of the perimeter of randomly chosen boutons and dendrites also revealed no significant differences between +/+ (boutons: 2.02 ± 0.57 µm, n = 75; dendrites: 2.58 ± 0.81 µm; n = 75) and -/- (boutons: 2.08 ± 0.83 µm; n = 75; dendrites: 2.46 ± 0.85 µm; n = 75) mice (n= 3 each; boutons: p = 0.967; dendrites: p = 0.389, Mann Whitney tests). However, we found a dramatic reduction of ∼ 50% in the total volume of astrocytic processes in the neuropil of the SON from -/- mice (n = 9, Fig. 2d-f), including around synaptic complexes, when compared with +/+ mice (n = 9, p = 0.0001, Mann Whitney test, Fig. 2d, f, g, i). We indeed observed a nearly 50% reduction in the percentage of astrocytic processes around synaptic boutons (n = 75 boutons) and dendrites (n = 75 dendrites) in -/-mice compared to +/+ mice (boutons: n = 75, p = 0.0009; dendrites: n = 75, p = 0.0005, Mann Whitney tests, Fig. 2d-i). Consistent with the reduced astrocyte volume and diminished synaptic coverage, we observed an increased distance by ∼ 65% between astrocytic processes and synapses (n = 120; p < 0.0001, Mann-Whitney test, Fig. 2i). In all, these results indicate that disruption of Cx30 expression decreases the astroglial coverage of OT-expressing neurons in the SON of virgin pup-naïve female mice.

**Fig. 2.**
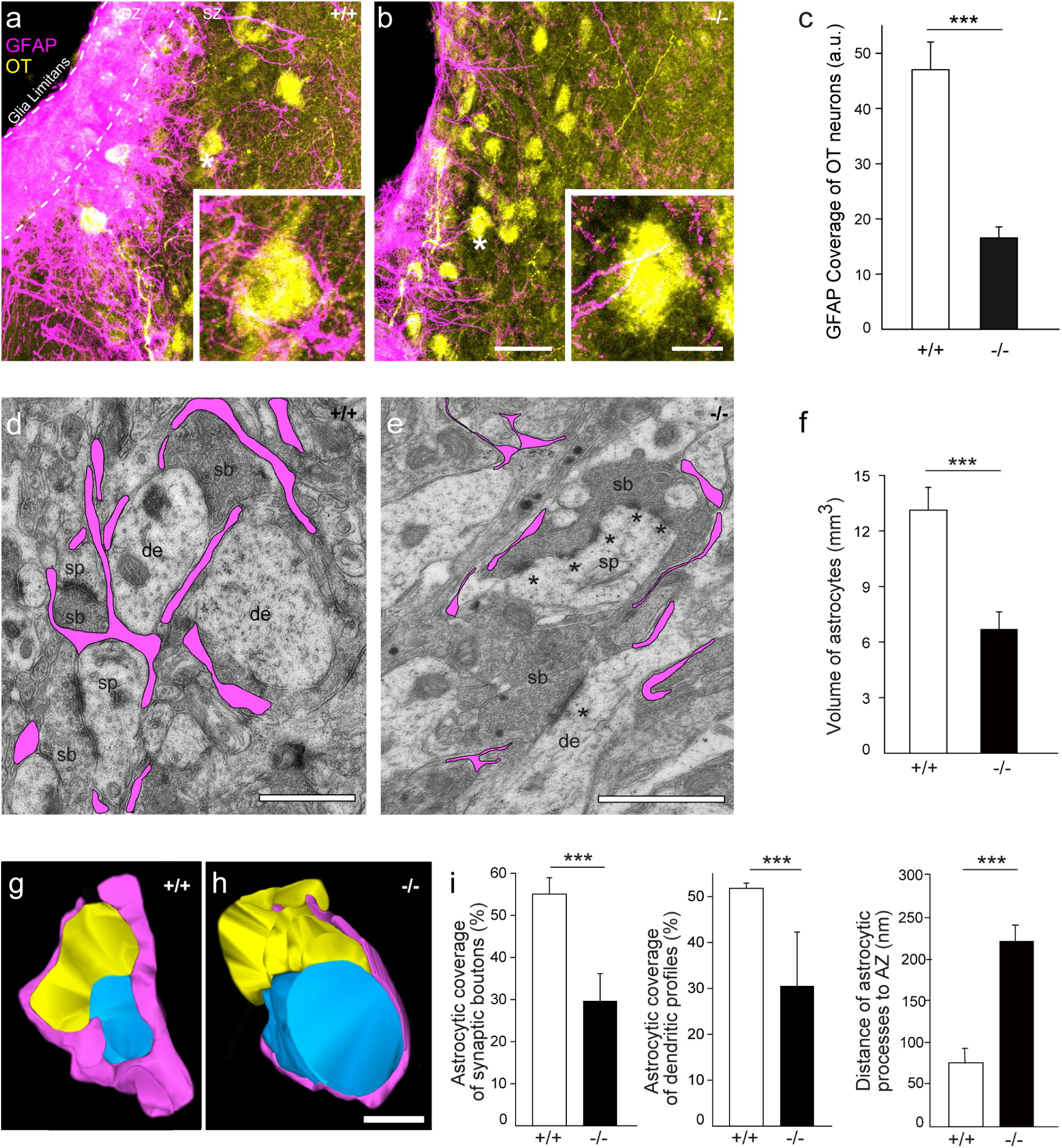
Cx30 regulates astroglial coverage of oxytocinergic neurons and synapses in the SON. Immunofluorescent images of GFAP (magenta) and OT (yellow) in the SON of +/+ (**a**) and -/- (**b**) mice. Representative oxytocinergic neurons covered by GFAP-rich astrocyte processes, as indicated by the white asterisks, are magnified in the insets. Scale bar: 30 µm; inset: 8 µm. DZ: Dendritic zone. SZ: Somatic zone. **c** Quantitative analysis of the GFAP coverage of OT-expressing neurons. -/- mice display reduced astroglial neuron coverage (n = 182) compared to control SON (n = 134). **d** High power electron micrograph of synaptic complexes from the SON of +/+ mice composed of either dendrites (de) or spines (sp) with terminating synaptic boutons (sb). Here, the fine astrocytic processes are highlighted in magenta. Scale bar: 0.5 µm. **e** High power electron micrograph from the SON of -/- mice showing two large synaptic boutons (sb), one terminating on a dendrite (de) and another on a spine (sp). Note several Active Zones (AZs) in the synaptic bouton on the spine (marked by asterisks) and the large, but non-perforated AZ at the dendrite. Scale bar: 1 µm. **f** Bar histograms showing a reduction in the volume of astrocytic processes (n = 9, p = 0.0001, Mann Whitney test) **g** 3D-volume reconstruction of a synaptic complex (yellow, synaptic bouton; blue, target spine) and the surrounding astrocytic processes (magenta) in the SON of +/+ mice. **h** Representative example of a synaptic complex and fine astrocytic processes in the SON of -/- mice. Note that astrocytic processes are less prominent and not as tight around the synaptic complex when compared with the +/+ mice. Same color code as in (**g**). Scale bar in (**g**) and (**h**): 0.5 µm. **i** Bar histograms showing a reduction of the percentage of astrocytic processes around synaptic boutons (n = 75) and dendrites (n = 75) and an increase in the distance of astrocytic processes to AZ in the SON of -/- mice compared to +/+ mice (% of astrocytes around boutons: n = 75, p = 0.0009; % of astrocytes around dendrites: n = 75, p = 0.0005, distance astrocytic process-AZ , n = 120; p < 0.0001, Mann Whitney tests). Asterisks indicate statistical significance (***p < 0.001).

### Cx30 deficiency increases plasma OT levels

Neuroglial structural plasticity in the SON has been shown to modulate the tone of magnocellular synaptic inputs, thereby promoting bursting activity and OT release in the bloodstream^28^. Hence, given the reduced astroglial synaptic coverage observed in the SON of virgin pup-naïve female -/- mice, we investigated whether Cx30 controls the plasma levels of OT. Using radioimmunoassay (RIA) (Fig. 3a), which allows specific quantification of basal OT levels^29^, we found that virgin pup naïve female -/- mice displayed a two-fold increase in plasma OT levels compared to wildtype virgin pup naïve female mice (+/+: 45.5 ± 2.5 pg/ml, n = 8, -/-: 87.6 ± 16.5 pg/ml, n= 6, p = 0.05, unpaired t-test, Fig. 3b). This indicates that astroglial Cx30 regulates OT plasma levels in virgin pup naïve female mice.

**Fig. 3.**
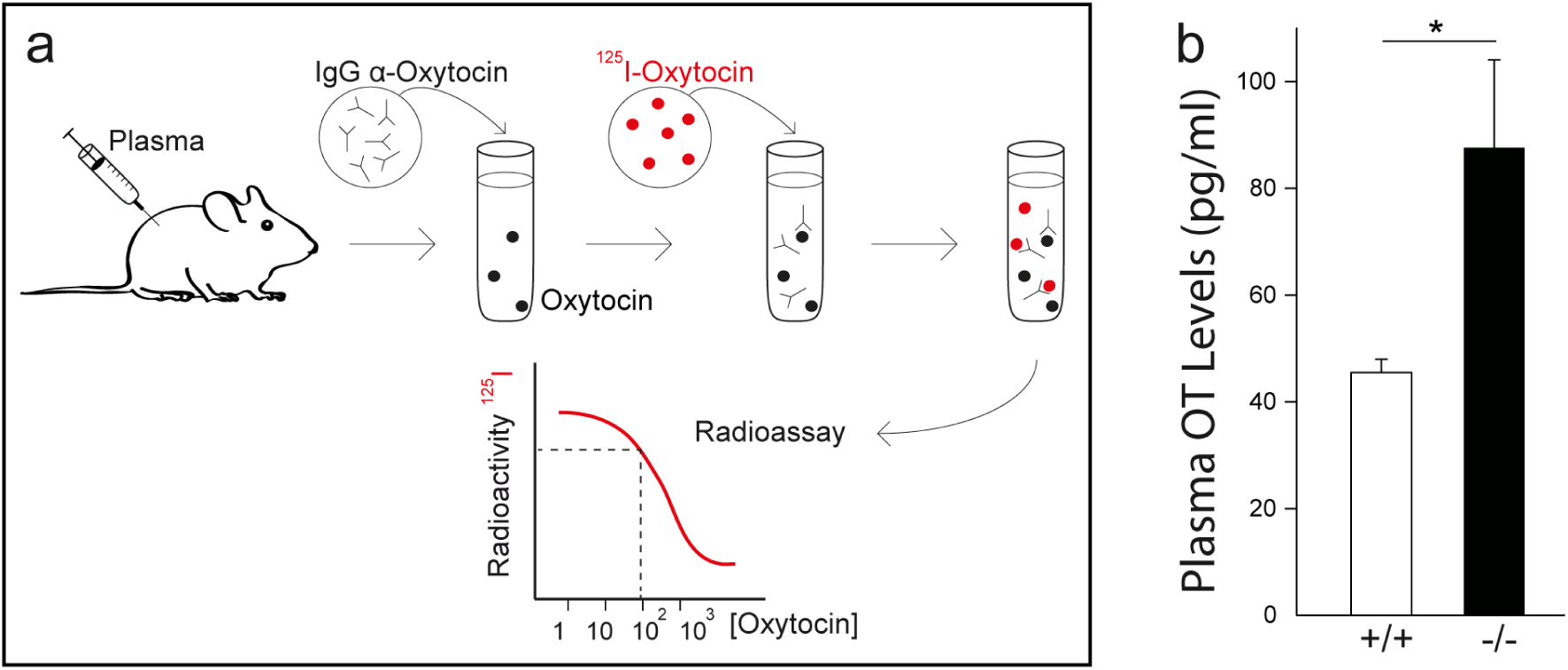
Cx30 sets plasma levels of OT. **a** Schematic representation of the radioimmunoassay (RIA) method used to measure OT plasma levels in mice. **b** Plasma OT levels measured by RIA were increased in -/- (n = 6) compared to +/+ mice (n = 8, p = 0.05, unpaired t-test). Asterisks indicate statistical significance (\**p =* 0.05).

### Cx30 deficiency promotes maternal behavior via regulation of OT signaling

OT is a prosocial neuropeptide having critical roles in affiliative behaviors, including especially maternal care^8^. Because virgin pup-naïve female mice deficient for Cx30 had increased plasma OT levels, we investigated whether they display enhanced maternal behavior. To this end, we first tested virgin pup-naïve females for maternal care abilities, which were assessed by quantifying their time spent in affiliative interaction with pups, including sniffing, nursing or moving the pups (Fig. 4a). We found an increased duration of female-pup affiliative interactions in -/- mice (n = 6) as compared to control (n = 8, p = 0.0068, two-way repeated measures ANOVA, Fig. 4b). We then tested virgin pup-naïve female mice for pup retrieval, an OT-dependent maternal behavior consisting in picking up an isolated pup emitting distress ultrasonic vocalizations and bringing it back to the nest^30^ (Fig. 4c). -/- mice (n = 9) showed significantly higher pup retrieval rates as compared to wildtype mice (n = 12, p = 0.015, two-way repeated measures ANOVA, Fig. 4d), suggesting higher maternal competences in female mice. Finally, pup-naïve female mice were also tested for nest building, a behavior involving the creation of a shelter from raw materials found in the environment to protect the pups^31^ (Fig. 4e). Virgin pup naïve female -/- mice (n = 9) displayed better nest capacities as compared to control (n = 10, p = 0.0083, two-way repeated measures ANOVA, Fig. 4f). We then tested whether this effect was sex specific. We found that -/- males (n = 8) displayed similar nest building capacity compared to +/+ males (n = 10, p = 0.627, two-way repeated measures ANOVA, Supplementary Fig. 4a, b), and that, as previously reported^32^, +/+ males (n = 10) and +/+ females (n = 10) had also comparable performances (p = 0.708, two-way repeated measures ANOVA, Supplementary Fig. 4c). These data indicate that the Cx30 regulation of nest building capacity was specific to female mice (-/- male: n = 8, -/- female: n = 9, p = 0.0147, two-way repeated measures ANOVA, Supplementary Fig. 4d). The increase in maternal care observed in -/- virgin female mice was specific with no effects detected on locomotion or anxiety (-/- n = 8, +/+ n = 6, p = 0.19 unpaired t-test and p = 0.35 respectively, Supplementary Fig. 5a-c). Altogether, these results indicate that -/- virgin pup-naïve female mice display exacerbated maternal behaviour.

**Fig. 4.**
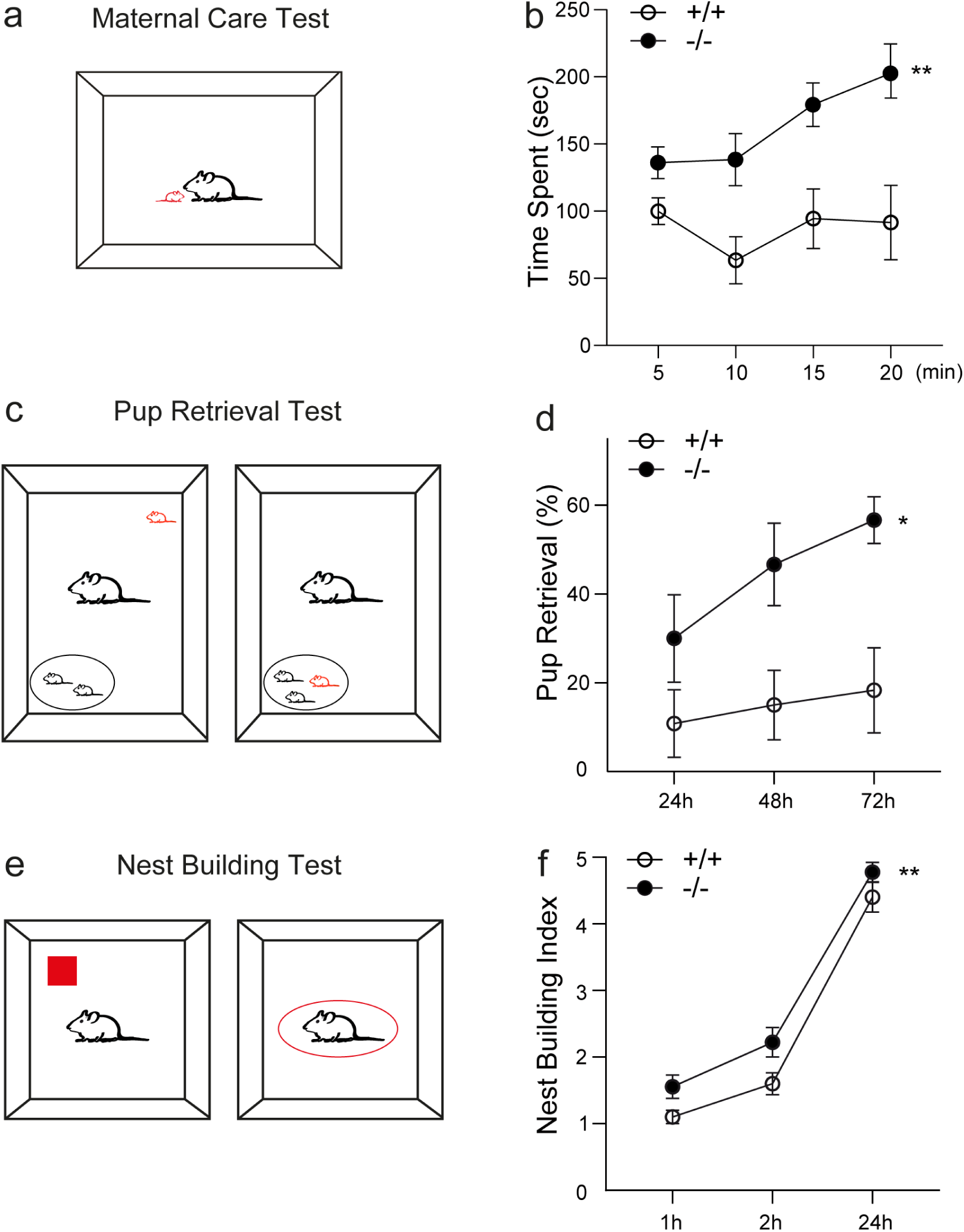
Cx30 modulates maternal behavior. **a** Schematic illustration of the maternal care test. **b** Virgin pup-naïve female -/- mice (n = 6) spent more time in affiliative interaction with pups (sniffing, nursing or moving the pup) compared to +/+ mice (n = 8, p = 0.0068, two-way repeated measures ANOVA). **c** Schematics representing the pup retrieval test. **d** Virgin pup naïve female -/- mice (n = 9) show increased retrieval rates as compared to +/+ (n = 12, p = 0.015, two-way repeated measures ANOVA). **e** Schematic illustration of the nest building test. **f** Virgin pup-naïve female -/- mice (n = 9) display improved nest building performances compared to +/+ (n = 10, p = 0.0083, two-way repeated measures ANOVA). Asterisks indicate statistical significance (\**p* < 0.05; \*\**p* < 0.01).

Since we showed increased plasma OT levels in virgin pup naïve female -/- mice, we then investigated whether Cx30 controls maternal behavior via activation of OT signaling. Virgin pup-naïve female -/- mice injected intraperitonally with an OT receptor (OTR) antagonist (L368 899, 10 mg/kg)^33^, which crosses the blood brain barrier, showed reduced scores on the three maternal behaviors that we assessed, i.e. maternal care (n = 7, p = 0.0412, two-way repeated measures ANOVA, Fig 5a b), pup retrieval (n = 7, p < 0.0001, two-way repeated measures ANOVA, Fig. 5c, d) and nest building tests (n = 7, p < 0.0001, two-way repeated measures ANOVA, Fig. 5e, f) compared to saline-injected virgin pup-naïve female -/- mice (n = 11, 6 and 7, respectively). In contrast, virgin pup-naïve female +/+ mice injected intraperitonally with the same OTR antagonist displayed similar poor maternal care (n = 10, p = 0.7212 two-way repeated measures ANOVA, Fig. 5a, b), pup retrieval (n = 7, p = 0.7134, two-way repeated measures ANOVA, Fig. 5c, d) and nest building performances (n = 9, p = 0.7361, two-way repeated measures ANOVA, Fig. 5e, f) compared to saline-injected virgin pup-naïve female +/+ controls (n = 9, 9 and 9, respectively). These data indicate that the maternal care behavior of virgin pup-naïve -/- mice is mediated by OT signaling. Interestingly, the changes in Cx30 and GFAP expression triggered by pup exposure were not affected by the injection of OTR antagonist in virgin +/+ mice (pup + saline: n = 21 ROI from 5 mice, pup + anta-OTR: n = 22 ROI from 5 mice, Cx30: p = 0.8955, unpaired t-test, GFAP: p = 0.5554, Mann Whitney test, Supplementary Fig 6a, b), suggesting that this process occurs prior to the increase in OT levels in the SON. Furthermore, in virgin -/- females previously repeatedly exposed to pups (pup-sensitized (n = 8) by 10 sessions of 2 minutes pup exposure daily during 3 days), which already experienced OT-dependent plasticity in the MPOA resulting in facilitation of maternal behavior^19^, the OTR antagonist had no effect on pup retrieval performance tested one month after the pup sensitization compared to -/- mice (n = 9, p = 0.702, two-way repeated measures ANOVA, Supplementary Fig. 7). This indicates that pup retrieval after maternal learning was independent of OT signaling in -/- mice, as previously reported in +/+ females^30^. Since the facilitation of maternal behavior is mediated by activation of the OT system in the MPOA^19^, we tested the implication of this circuit. To do so, we stereotaxically injected an OTR antagonist (d(CH2)51,Tyr(Me)2,Thr4,Orn8,Tyr-NH29)-vasotocin trifluoroacetate salt) in the MPOA of virgin pup-naïve female -/- mice (n = 7). The OTR antagonist decreased the maternal care abilities of these mice compared to saline controls (n = 6), as shown by the amount of time spent in interaction with pups (p = 0.0155, two-way repeated measures ANOVA, Supplementary Fig. 8a, b). In contrast, virgin pup-naïve female +/+ mice locally injected in the MPOA with the same OTR antagonist (n = 8) did not show altered female-pup interaction duration when compared to saline control (n = 8, p = 0.3913, two-way repeated measures ANOVA, Supplementary Fig. 8a, b). Altogether, these data indicate that astroglial Cx30 deficiency exacerbates maternal behavior via activation of the oxytocinergic system in the MPOA.

**Fig. 5.**
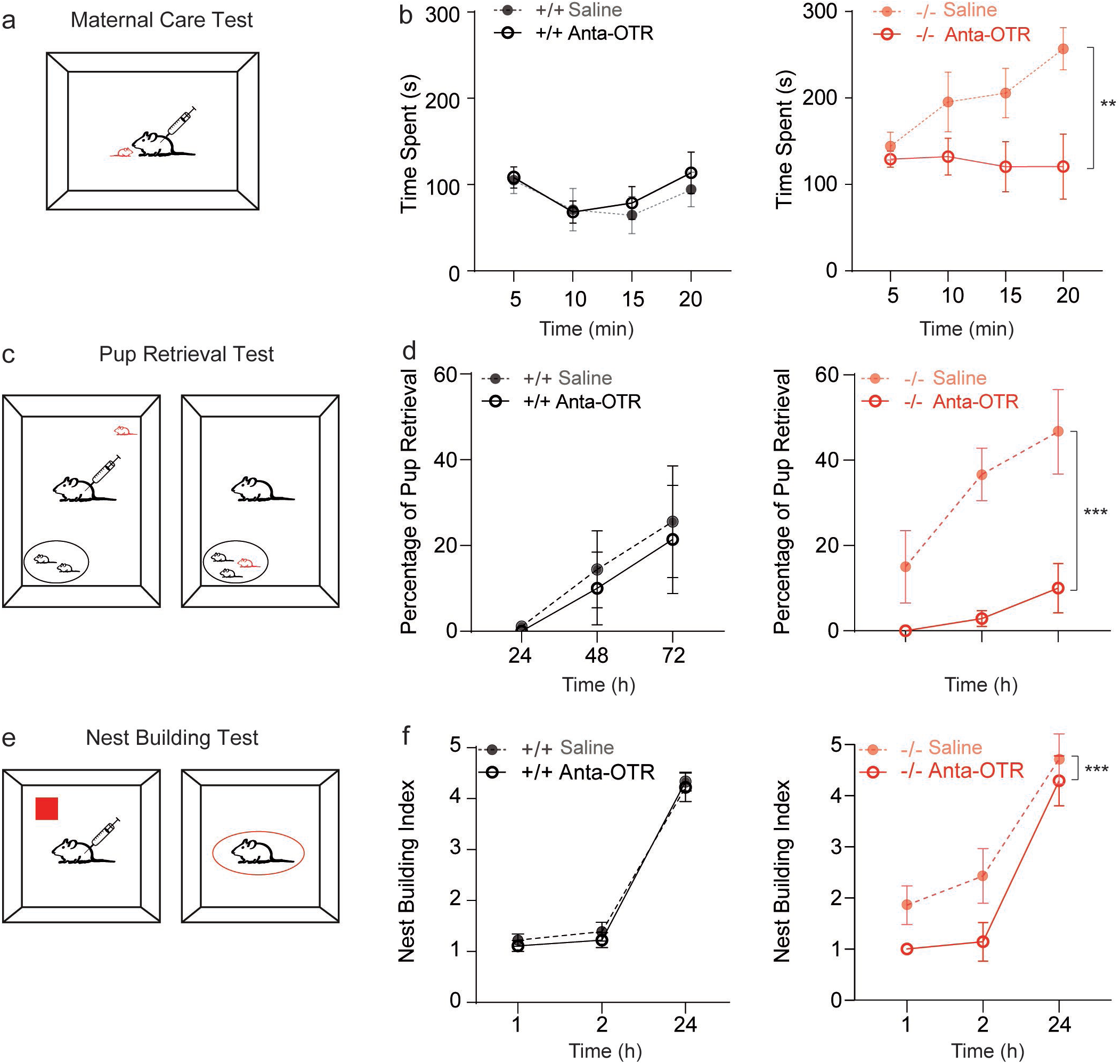
Cx30 controls maternal behaviour through OT signaling. Schematic representation showing the maternal care (**a**) pup retrieval (**c**) and nest building (**e**) procedures used to assess pup-directed and indirect maternal care in +/+ and -/- mice injected intraperitoneally with saline or the blood brain barrier permeable OTR antagonist (Anta-OTR, L368 899) 30 minutes before the test. Maternal care (**b**), pup retrieval (**d**) and nest building (**f**) performances of +/+ mice injected with the L368 899 (n = 10, 7 and 9, respectively) were similar to the ones of +/+ mice (n = 9, p = 0.7212; n = 9, p = 0.7134 and n = 9, p = 0.7361, respectively, two-way repeated measures ANOVA). In contrast, performances of -/- mice injected with the L368 899 (n = 7, 7 and 7, respectively) differs significantly from -/- mice (n = 11 p = 0.0084, n = 6, p < 0.0001 and n = 7, p <0.0001, respectively, two-way repeated measures ANOVA), thus indicating that Cx30 controls maternal behaviour via OT signaling. Asterisks indicate statistical significance (\*\**p* < 0.01; ***p < 0.001).

### Astroglial Cx30 in the extended SON regulates plasma OT levels and maternal behavior independently of its channel function

We then investigated whether disrupted astroglial Cx30 expression in the SON can increase plasma OT levels and promote associated maternal behaviurs. To this end, we delivered adeno-associated virus serotype 2/9 (AAV2/9) to express either GFP-tagged Cx30 (AAV2/9 Cx30-GFP) or GFP (AAV2/9 GFP) only as a control, under the GFAP promoter (gfaABC1D), in the SON of virgin pup naïve -/- or +/+ female mice, respectively (Fig. 6a and Supplementary Fig. 9). This resulted in hypothalamic expression of GFP in the SON and close adjacent areas, here called extended SON, including the lateral hypothalamic area, retrochiasmatic area and anterior hypothalamic nucleus (Supplementary Fig. 9a). Local viral transduction of transgenes under the GFAP promoter also reached SON close neighboring regions, as the SON is a very small structure, and there is no known SON specific marker of astrocytes to restrict astroglial targeting in this region. The specificity of the viral transduction in astrocytes was tested by co-immunolabeling of GFP, GFAP and Cx30 (Fig. 6). We found that GFP+ cells expressed GFAP, and that restoration of Cx30 expression occurs in GFP and GFAP expressing cells, confirming specific viral targeting of astrocytes. In addition, single cell analysis in the SON of virgin pup naïve -/- female mice transduced with AAV2/9 Cx30-GFP in astrocytes revealed that Cx30 expression in GFP-positive (GFP+) cells was increased by ∼25%, as compared to injection of AAV2/9 GFP in virgin pup naïve +/+ female mice, indicating restoration of Cx30 levels in -/- mice (Supplementary Fig. 9a, b).

**Fig. 6.** Astroglial Cx30 in the SON regulates plasma OT levels and maternal behaviour. **a** Diagram representing the strategy to restore Cx30 expression levels in the SON of -/- mice using local stereotaxic injection of astrocyte-targeted adenoviral vectors encoding GFP and Cx30 (AAV Cx30-GFP). OB: olfactory bulb, SON: supraoptic nucleus, PVN: paraventricular nucleus, CB: cerebellum. **b** GFP immunofluorescence labeling (green) in the SON of -/- mice locally injected with AAV encoding Cx30. Scale bar: 100 µm. **c** Immunofluorescence images showing restoration of Cx30 expression (yellow) in a GFP- (green) and GFAP-expressing (magenta) astrocyte from the SON of -/- mice. Scale bar: 20 µm. **d** Schematic representation of the radioimmunoassay method used to measure OT plasma levels in mice. **e** Normalized plasma OT levels in virgin pup naïve female -/- mice locally infected by AAV encoding Cx30 and GFP (light green, n = 6) were reduced compared to virgin pup naïve female -/- mice infected by control AAV encoding (dark green, n = 14, p = 0.0382, unpaired t-test). Schematic representation showing the maternal care (**f**) pup retrieval (**h**) and nest building (**j**) procedures used to assess pup-directed and indirect maternal care in virgin pup naïve female -/- mice locally infected by AAV encoding Cx30 and GFP or GFP only. Maternal care (**g**), pup retrieval (**i**) and nest building (**k**) performances of virgin pup naïve female -/-mice locally infected by AAV encoding Cx30 and GFP (n = 6, 8 and 10, respectively) were significantly reduced compared to virgin pup naïve female -/- mice locally infected by control AAV encoding GFP only (n = 10, p=0.0035; n=8 p=0.0039 and n=8, p=0.0016, respectively two-way repeated measures ANOVA). Asterisks indicate statistical significance (*p < 0.05; **p < 0.01; ***p < 0.001).

Restoring *in vivo* Cx30 expression selectively in astrocytes of virgin pup-naïve female -/-mice, using bilateral injection of adenoviral vectors encoding Cx30-GFP in the SON (AAV Cx30-GFP, Fig. 6a-c and Supplementary Fig. 9a,b), decreased OT levels by ≍ 40 % compared to virgin pup naïve -/- + GFP female mice (normalized OT levels to +/+ + GFP: -/- + Cx30-GFP: 1.07 ± 0.19, n = 6, -/- + GFP: 1.817± 0.21, n = 13, p = 0.0382, unpaired t-test, Fig. 6d, e), reinstating wildtype values. Restoring Cx30 expression in astrocytes significantly enhanced the expression of GFAP and the coverage of neuron somas by astrocytic processes (+/+ + GFP: n = 22 astrocytes from 10 slices from 6 mice, -/- + Cx30-GFP: n = 25 astrocytes from 20 slices from 6 mice, p = 0.0113 (GFAP) and p < 0.0001 (neuron coverage), unpaired t-test, Supplementary Fig. 9c-f). Consistently, restoring Cx30 expression in -/-mice significantly decreased the maternal performances: reduction of the time spent with pup in maternal care test (-/- + Cx30-GFP: n = 6, -/- + GFP: n = 10, p = 0.0035, two-way repeated measures ANOVA, Fig. 6f, g), decrease in pup retrieval rate (-/- + Cx30-GFP: n = 8, -/- + GFP: n = 8, p = 0.0039, two-way repeated measures ANOVA, Fig. 6h, i), and quality of nest building (-/- + Cx30-GFP: n = 10, -/- + GFP: n = 8, p = 0.0016, two-way repeated measures ANOVA, Fig. 6j-k). These data demonstrate that astroglial Cx30 in the extended SON regulates plasma OT levels and maternal behavior.

Besides its well-known role in GJ-mediated intercellular coupling, Cx30 also forms hemichannels, allowing direct exchange with the extracellular space, as well as non-channel functions such as adhesion or protein interactions^34^. We thus investigated which function of Cx30, channel or non-channel, mediates the regulation of plasma OT levels and associated maternal behavior in virgin pup-naïve female mice. To this aim, we used the Cx30T5M mice that express a mutated form of Cx30 in which the replacement of a threonine by a methionine at position 5 inhibits intercellular biochemical coupling mediated by GJ channels^35^. In these mice, plasma OT levels and nest building performances were similar compared to virgin pup-naïve female +/+ mice and significantly differed from -/- mice (Normalized plasma OT levels to wildtype: -/-: 1.93 ± 0.36, n = 6, Cx30T5M: 1.06 ± 0.13, n = 9, , p = 0.02 unpaired t-test, Supplementary Fig. 10a, b; Nest Building Index: -/- n = 9, Cx30T5M: n = 8, p = 0.007, two-way repeated measures ANOVA, Supplementary Fig. 10c, d). Accordingly, the quality of nest building in Cx30T5M mice significantly differed from that of -/- mice (Cx30T5M: n = 8, -/-: n = 9, p = 0.0133, two-way repeated measures ANOVA, Supplementary Fig. 10d). Altogether, these results indicate that astroglial Cx30 in the SON regulates plasma OT levels and maternal behavior, independently of GJ-mediated biochemical coupling.

## DISCUSSION

The present study shows that in a situation of social experience-dependent structural plasticity associated with increased OT release, virgin female mice display downregulation of astroglial Cx30 in the hypothalamic SON. Moreover, we found that in a mouse model already disrupted for Cx30, astroglial coverage of oxytocinergic neurons and synapses is decreased in the SON of virgin females. This translated at the functional level in elevated plasma OT levels and at the behavioral level in exacerbation of maternal care by activation of OT signaling in the MPOA. Finally, we showed that astroglial Cx30 in the extended SON set plasma OT levels and maternal behavior, independently of GJ-mediated biochemical coupling. Altogether these data establish hypothalamic astrocytes, via Cx30, as key regulators of OT-based maternal behavior.

### Astrocytes as regulators of OT-dependent social behaviors

Alteration of astrocyte-specific functions has recently been held accountable for impairments in social cognition and behavior^36, 37^. In addition, findings from postmortem brain studies and animal models of psychiatric disorders characterized by social deficits have unraveled clinical signs of impaired astrocytes’ physiology^38, 39^. However, so far, there was no direct evidence of astroglial regulation of neuroendocrine neuronal circuits involved in social interactions. Here we show that in virgin pup-naïve female mice, pup exposure transiently decreases in the SON the expression of Cx30, which is associated with astrocyte structural remodeling, consisting in reduced morphological complexity and OT neurons coverage, and increased OT levels. Remarkably, we found that Cx30 expression regulates the structural properties of astrocytes and OT circuitry controlling maternal behavior. Indeed, Cx30 deficiency in virgin pup-naïve female mice reduces local astrocyte ensheathment of OT neurons in the SON and increased plasmatic OT levels, which activates the MPOA circuitry, the central fulcrum for processing pup-related cues^20^, exacerbating maternal behavior. It is noteworthy that virgin pup naïve female wildtype mice display low levels of basal plasmatic OT and poor maternal behavior^30^, as we here show with low pup retrieval success rate (∼10%), pup interaction time and nest building performance compared to -/- mice (∼ -50%). Because OTR are likely weakly activated in wildtype virgin female mice, their inhibition has no significant effect on maternal behavior driven by OT signaling.

The Cx30-mediated rise in plasma OT is likely accompanied by elevated levels of OT release within the SON as well as within OT neuronal target regions, as seen during suckling^40, 41^ or chemogenetic stimulation of OT neurons^42^. Such intracerebrally released OT has been shown to be essential for various social behaviors including maternal behavior^43^.

Remarkably, OTR are expressed in MPOA neurons and up-regulated perinatally, most likely in response to pup-derived stimuli^44^. Pup exposure increases OT levels within the MPOA^19^, and induces cFos immediate early gene enrichment in MPOA OTR-expressing neurons of pup-sensitized females^19^ and dams^45^. Moreover, activation of MPOA OTR elicits maternal responsiveness in both pup-sensitized females^19^ and dams^46^. Thus, the astroglial Cx30-mediated activation of MPOA OTR must play an instructive role in the maternal behavioral transition that female mice undergo following exposure to pups.

Astrocytes express most, if not all, receptors for neuroactive molecules, and are thus able to mediate some of their neuromodulatory actions^47^. The astroglial molecular repertoire includes notably behavior-related neurohormone receptors, such as dopamine^48^ and opioid^49^ receptors, whose physiological roles have only begun to emerge. In particular, expression of OTR in astrocytes has been reported in various brain areas^50, 51^, including the SON^52^. Moreover, OT has recently been shown to regulate emotional and social behaviors not only through direct modulation of neuronal networks, but also via activation of OTR expressed on astrocytes. OT indeed acts on astrocytes of the central amygdala, a brain region associated with emotional processing, to regulate neuronal excitability through release of the NMDAR co-agonist D-serine, thereby modulating behavioral fear responses and associated anxiety levels^51^. OTR signaling in astrocytes from different brain regions, by modulating local neuronal activity, may thus contribute to distinct behavioral responses.

### Astroglial coverage of synapses as a mechanism to regulate physiological functions

Experience-dependent brain plasticity not only occurs at synapses, but also takes place at the point of contact between neurons and glial cells, including in particular astrocytes^53^. In line with this, we here report that SON astroglial Cx30 protein levels, independently of OT signaling, are dynamically downregulated following social experience-dependent plasticity, and that Cx30 deficiency leads to a reduction in astroglial volume and coverage of oxytocinergic neurons in the SON, controlling OT levels. These results suggest the following sequence of events: pup exposure, within 30 minutes, decreases Cx30 expression, which induces astrocyte structural remodeling consisting in reduced GFAP levels and morphological complexity, resulting in decreased astroglial coverage of OT neurons at the somatic level and increased OT levels. In this scenario, the morphological changes precede the increase in OT. These alterations are dynamic, as they reversed 90 min after pup exposure, except for the astroglial coverage of OT soma, which remains reduced at this timepoint. It is however possible that 90 min after pup exposure, it is at the synaptic level that this astroglial coverage is increased and thus reversed, which limits excitatory synaptic activity via enhanced glutamate clearance and reduced spillover, and thereby decreases OT release at the population level, restoring OT levels back to baseline. Alternatively, one can hypothesize that the increase in astroglial morphological complexity is sufficient to decrease OT levels back to baseline 90 min after pup exposure, via an overall decrease in extracellular space volume, which can also result in reduced glutamate spillover.

Noteworthy, unlike in cortical regions, physiological neuroglial remodeling in the SON leads to dramatic changes in astroglial ensheathment of synapses, which strongly impacts neuronal function through various mechanisms. Previous studies have indeed shown that SON astrocyte-synapse proximity regulates, inter alia, extracellular space volume and geometry^54^, astroglial uptake of neurotransmitters^3^, release of astrocytic gliotransmitters^55^ and contact-mediated astrocyte-synapse signaling^56, 57^. With regard to the latter, astrocyte-neuron contacted-mediated signaling involves remodeling of the extracellular matrix. Remarkably, enzymes degrading the extracellular matrix are expressed in astrocytes and OT neurons of the SON. Their expression increases in physiological situations of high hormonal demand and are suggested to be involved in SON structural plasticity^58, 59^. Interestingly, astroglial Cx can regulate the extracellular matrix by controlling the expression and activity of matrix degrading enzymes^26, 60^, which may thus participate to the socially-induced regulation of neuroglial structural plasticity in the SON.

At a more integrated level, we here show that Cx30-dependent remodeling of SON astroglial synapse coverage translates into functional changes of OT circuits controlling social behavior, which likely result from regulation of synaptic activity. We have indeed previously shown that deficiency for Cx30 alters neuronal activity in brain areas such as the hippocampus or visual cortex^25, 26, 61, 62^, and that local hippocampal synaptic activity is directly regulated by astroglial coverage of synapses^25^. The mechanism proposed in our study is thus unlikely to result from compensatory alterations in -/- mice. Our data indeed point to the specific implication of astroglial Cx30 from the extended SON in the morphological and behavioral effects that we here report, as we show that restoring Cx30 expression selectively in astrocytes from this brain region rescues the structural and functional alterations that we found in -/- virgin female mice, namely GFAP levels, astroglial coverage of oxytocinergic neurons, oxytocin levels and maternal behavior. In addition, we successfully rescued wildtype maternal behavior in -/- mice by acutely administering an oxytocin receptor antagonist.

Remarkably, a few other studies point to a contribution of astroglial morphological remodeling to cognitive and behaviural processes during physiological states or diseases. For instance, the astrocytic leptin receptor-mediated remodeling of astroglial neuron coverage in the hypothalamic arcuate nucleus regulates the synaptic input of neuronal feeding circuits, namely pro-opiomelanocortin (POMC)- and Agouti-related protein (AgRP)-producing neurons, and hence control feeding behaviors^63^. Therefore, astrocytic synapse ensheathment emerges as an important mechanism of astrocyte neuromodulation. Interestingly, astrocyte synapse coverage dynamically changes with synaptic activity and plasticity, and appears to be region-specific. For example, whereas in the hippocampus induction of long-term potentiation (LTP) causes perisynaptic astrocyte processes to approach activated synapses^64, 65^, in the amygdala, fear learning induces the reduction of astrocytic physical contacts with potentiated synapses^66^. These differences in activity-dependent astroglial synapse coverage may originate from region-dependent heterogeneity of astrocytes and result in regulation of specific circuits underlying behavior. It is noteworthy that the Cx30 regulation of astroglial synapse coverage varies according to the brain area, as in Cx30 deficient mice, the coverage is increased in the hippocampus^25^, whereas we here show that it is decreased in the SON. Our recent data indicate that Cx30 is a molecular brake for structural plasticity in astrocytes, by controlling the actin cytoskeleton and mechanical remodeling^67^. Therefore, the reduction in Cx30 expression is permissive for astroglial structural plasticity, but does not determine the direction of the plasticity, i.e. retraction or extension of perisynaptic astroglial processes. This is likely controlled by other specific local factors, such as neurotransmitters or neuromodulators, which may underlie the brain-region heterogeneity of astrocytes^68^.

Altogether, these data demonstrate that astrocytes, via Cx30, actively participate in shaping social behavior mediated by neuroendocrine circuits, and may shed light on the etiology of mental disorders. This may contribute to genetic disorders associated with hypersociability, such as the Williams syndrom, a neurodevelopmental disorder characterized by increased OT plasma levels^69^. Furthermore, excessive maternal care can generate anxiety and depression, impair cognition and emotional behavior, inducing social dysfunction in adulthood in rodents and humans^70–72^. As a matter of fact, Cx30 is impaired in the brain of animal models of major depressive disorder^73, 74^, as well as patients with major depression^75^ and suicide completers^76^.

## Supporting information

Supplementary informations

## ACKNOWLEDGEMENTS

This work was supported by grants from the European Research Council (Consolidator grant #683154), the European Union’s Horizon 2020 research and innovation program (Marie Sklodowska-Curie Innovative Training Networks, grant #722053, EU-GliaPhD), the ANR under France 2030 (Chaire d’excellence SensoPAP, ANR-24-CHBS-0007), the Grand Prize by the Sicard Foundation-French Academy of Sciences, and the Major Research Program of PSL Research University “PSL-Neuro” launched by PSL Research University and implemented by ANR (ANR-10-IDEX-0001) to N.R., the Sorbonne University doctoral school ED3C to G.G. and M.G., Labex Memolife to G.G., and the German Research Council (Graduate School 2174) to I.D.N. We thank M. Cohen-Salmon for providing the -/- mice, and all members of the Neuroglial Interactions in Cerebral Physiology and Pathologies laboratory for helpful discussions. In addition, we would like to thank Brigitte Marshallsay and Ulrike Holz for excellent technical assistance in the electron microscopic part of the study. Furthermore, we would like to thank Dr. Kurt Saetzler for statistical advice.

## AUTHOR CONTRIBUTIONS

Conception and experimental design: A.M., C.L.C., I.A., G.G., J.L., J.M., and N.R.; collection, analysis and interpretation of the data: A.L., A.R., C.L.C., C.M., G.G., I.A., I.N. J.L., J.M., M.G., P.B., V.C., N.M.P., P.E. and N.R.; Manuscript writing: I.A., J.M., G.G., M.G., J.L. and N.R.

