## Supplementary informations for "Astrocytes control oxytocin-based maternal behavior via connexin 30"

**This file includes**

**Supplementary Materials and Methods**

**Supplementary Figures 1-10**

### Supplementary Materials and Methods

#### Animals

Experiments were carried out according to the guidelines of the European Community Council Directives of January 1st 2013 (2010/63/EU) and of the local animal welfare committee (certificate A751901, Ministère de l'Agriculture et de la Pêche), and all efforts were made to minimize the number of animals used and their suffering.

Experiments were carried out using adult virgin pup-naïve female mice of wild-type C57BL/6j background (+/+), GFAP-eGFP mice[1], in which the enhanced green fluorescent protein (eGFP) is expressed under the GFAP promoter, knockout mice for Cx30 (-/-)[2] and Cx30T5M[3] mice, unless otherwise stated. These mice were generated and characterized as previously described[1-3]. GFAP-eGFP mice were provided by H. Kettenmann (Max-Delbrück Center for Molecular Medicine, Berlin, Germany) and Cx30T5M mice were provided by F. Mammano (Venetian Institute of Molecular Medicine, Italy).

Animals were group housed on a 12 h light/dark cycle. All mice were backcrossed to the C57BL/6j background. Experiments were performed on adult female mice (2-4 months), unless otherwise stated.

#### AAV production and stereotaxic injection in the SON

For AAV in vivo gene transfer, a transgene composed of GFP and Cx30 cDNA separated by a P2A sequence in a single open reading frame was placed under the control of a GFAP-specific promoter in an AAV shuttle plasmid containing the inverted terminal repeats (ITR) of AAV2. Pseudotyped serotype 9 AAV particles were produced by transient co-transfection of HEK-293T cells, as previously described[4]. Viral titers were determined by quantitative

PCR amplification of the ITR on DNase-resistant particles and expressed as vector genome per ml (vg/ml).

Mice (age 3–4 weeks) were anesthetized with a mixture of ketamine (95 mg/kg; Merial) and xylazine (10 mg/kg; Bayer) in 0.9% NaCl and placed on a stereotaxic frame under body temperature monitoring. Buprenomorphine analgesics (0.1 mg/kg; Centravet) were then sub-cutaneously administered to the mice. AAVs were diluted in PBS with 1% BSA at a concentration of $0.5 \times 10^{13}$ vg/ml, and 1 µl of virus was stereotaxically injected bilaterally above the SON at a rate of 0.2 µl/min, using a 29-gauge blunt-tip needle linked to a 2 µl Hamilton syringe (Phymep). The corresponding stereotaxic coordinates were, from bregma: antero-posterior −0.57 mm, medio-lateral ± 1.05 mm and dorso-ventral −5.2 mm. At the end of the injection, the needle was left in place for 5 min before being slowly removed. The skin was sutured and mice were allowed to recover for at least 3 weeks before experiments were performed.

**Stereotaxic injection of OTR antagonist in the MPOA**

After intraperitoneal injection with an anesthetic mixture of ketamine (95 mg/kg; Merial) and xylazine (10 mg/kg; Bayer) in saline and subcutaneous administration of the buprenomorphine analgesic (0.1 mg/kg; Centravet), +/+ vs -/- female mice were kept warm using a heating pad and stereotaxically implanted with a 26-gauge bilateral guide cannula (3.35 mm long below pedestal, Plastic One) positioned above the MPOA (using the following stereotaxic coordinates, from the bregma : antero-posterior -0.14 mm, medio-lateral ±0.6 mm, and dorso-ventral -2.7 mm). The canula was held in place by dental cement. After surgery, mice were re-housed in their home cage to recover for at least 7 days under close monitoring. After the recovery period, and 20 minutes before behavioral testing, animals were bilaterally injected in the MPOA with 0.1 µl of either saline or an OTR antagonist (10 mg/kg, (d(CH2)51,Tyr(Me)2,Thr4,Orn8,Tyr-NH29)-vasotocin trifluoroacetate salt, Fisher Scientific) dissolved in saline at a concentration of 0.5 µg/µl. This was achieved through a 33-gauge bilateral canula, extending 2 mm below the end of the guide canula to target the MPOA, and linked to two 2 µl Hamilton syringes (Phymep) controlled by a syringe pump (KD Scientific), allowing injections at the rate of 0.1 µl/min.

**Antibodies, immunohistochemistry and immunoblotting**

All the antibodies used in this study are commercially available and have been validated in previous studies, as reported by the suppliers. The following primary antibodies were used for immunohistochemistry: GFAP mouse monoclonal (1:1000, G3893, Sigma-Aldrich) or GFAP rabbit polyclonal (1:500, G9269, Sigma-Aldrich), OT rabbit polyclonal (1:10,000, AB911, Merck Millipore) or OT guinea pig polyclonal (1:500, 408004, Synaptic System), Cx30 polyclonal rabbit (1:500, 71-2200, ThermoFisher) and GFP polyclonal chicken (1:500, 1020, Aves). The following fluorescent dye-conjugated secondary antibodies were used in appropriate combinations: goat anti-mouse IgG conjugated to Alexa 555 (1:2,000, A-21422, ThermoFisher), goat anti-rabbit IgG conjugated to Alexa 488 (1:2,000, A-11034, ThermoFisher), goat anti-mouse IgG conjugated to Alexa 647 (1:1000, A-21235, ThermoFisher), goat anti-rabbit IgG conjugated to Alexa 555 (1:1000, A-27039, ThermoFisher), goat anti-chicken IgG conjugated to Alexa 488 (1:2000, A11039, ThermoFisher), goat anti-rabbit IgG conjugated to Alexa 633 (1:2000, A21105, ThermoFisher).

Immunohistochemistry was performed as previously described[5]. Briefly, animals anesthetized with lethal dose of Euthasol (100µl /10g), were perfused by intracardiac with PBS first and 2% paraformaldehyde (PFA). The brains were carefully removed for an overnight post-fixation in the same fixative, followed by transfer in 30% sucrose for cryoprotection. Brain coronal microtome sections (40µm thick) were collected in PBS and pre-incubated 1 h with PBS-1% gelatin in the presence of 0.25% Triton-X100 (PGT). Brain sections were then stained overnight at 4°C with primary antibodies and washed in PGT three times. Appropriate secondary antibodies with DAPI (1:200, D9564, Sigma-Aldrich) were finally applied for 2 hours at room temperature. After several washes in PBS, brain slices were mounted in Fluoromount (Clinisciences) and examined with a spinning-disk X1 (CSUX1-A1, Yokogawa). Stacks of images were taken with a 63x immersive objective at 0.2 µm intervals.

Cell surface protein biotinylation and immunoblotting were performed using an adapted version of our previously described protocol [6]. Acute prefrontal cortex slices were prepared from control and pup-exposed WT female mice (2-3 months old) using a brain matrice (RBMS-205C, WPI), immediately after the behavioral test, and maintained on ice in artificial cerebrospinal fluid (aCSF). Slices were incubated for 1h on ice in a solution containing 1mg/mL biotine (EZ-Link® Sulpho-NHS-SS-Biotin, 21331, Thermo Scientific). Slices were washed three times in ice-cold aCSF and excess biotin was quenched by incubated the slices two times 25 min in cold quench buffer containing 100 mM glycine in aCSF. After three washes in ice-cold aCSF, slices were frozen in liquid nitrogen and the SON region was microdissected using a 22-gauge blunt needle (LS22 Phymep).

For each biological replicate, bilateral SON punches from 4 mice (one slice per mouse) were pooled to obtain sufficient material, therefore each sample contained tissue from 8 SON (4 mice x 2 SON). Samples were homogenized in the RIPA buffer containing: 100 mM Tris, 150 mM NaCl, 1 mM EDTA, 1% Triton-1X-100, 0.1% SDS and 1% Na deoxycholate supplemented with protease inhibitors cocktail and incubated for 30 min at 4°C to complete the lysis. Lysates were centrifuged at 16,000 x g for 15 min at 4°C, and supernatants were stored at -80°C until use. Protein concentration was determined using Ionic Detergent Compatibility Reagent (22663, Thermofisher).

For cell surface protein isolation, 60 µg of each sample was incubated with 20 µl of precleared streptavidin agarose beads (Pierce™ Streptavidin Plus UltraLink™ Resin, 53116, Themo Scientific) overnight at +4 °C. After centrifugation at 16,000 × g for 2 min at RT, the beads were washed three times in RIPA buffer and resuspended in 30 µl of 2× Laemli buffer. Samples were rotated for 1h at RT and then spun down. Twenty microliters of each biotinylated fraction were loaded on precast 4%–12% gradient gel (NuPAGE Novex Bis-Tris gel, NP0321BOX, Invitrogen). The proteins were transferred onto nitrocellulose membrane and saturated with 5% fat-free dried milk in triphosphate buffer solution. Membranes were incubated with primary antibody rabbit anti-Cx30 (1:1000, 71-2200, Thermofisher) or mouse anti-GFAP (1:1000, G3893, Sigma-Aldrich) at 4 °C overnight. The next day, membranes were incubated with horseradish peroxidase (HRP)-conjugated secondary antibody goat anti-rabbit-HRP (1:2000, CSA2115, Cohesion Biosciences) or goat anti-mouse-HRP (1:2000, CSA2108, Cohesion Biosciences) for 2 h at RT. Monoclonal mouse anti-β actin-HRP antibody (1:2000, clone AC-15, ab49900, Abcam) was used as a loading control. Protein bands were visualized using a chemiluminescence detection kit (ECL, 28980926, GE Healthcare) and imaged on an ImageQuant LAS 4000 system (GE Healthcare/Fujifilm).

Band intensities were quantified using ImageJ (NIH). For each lane, the region of interest corresponding to the target protein band was manually defined, and the area under the curve (AUC) of the pixel intensity profile was calculated.

**Imaging analysis**

*Cx30 and GFAP intensity in the SON.* Cx30 and GFAP integrated densities were measured in Z-projected images of the SON using ImageJ software. We normalized the data to the integrated density of the corresponding DAPI integrated density.

*GFAP coverage of OT neurons in the SON.* OT-labeled magnocellular somata were segmented using Otsu's thresholding algorithm on maximum intensity Z-projected images of the SON using ImageJ software. The GFAP fluorescent mean signal within OT-labeled magnocellular somata was then measured in maximum intensity Z-projected images of the SON.

*Three-dimensional analysis of astrocytes morphology.* Z-stack images were acquired at a spinning disk confocal microscope with a 60X magnification (CSUX1-A1 Yokogawa). Isolated astrocytes were selected as regions of interest based on their GFP staining in the SON of GFAP-eGFP mice. We used Imaris software (Oxford Instruments) to reconstruct astrocytes by employing the module filament tracer. All astrocytes with incomplete somata (cut in the x, y or z plane) were manually removed and not included in the further analysis. The following custom settings were used: largest diameter 7.00 µm, seed points 0.300 µm, remove seed points around starting points: diameter of sphere regions: 10.8 µm. Seed points were manually corrected (either placed in or removed from the center of the somata) if the IMARIS algorithm placed them incorrectly. All filament parameters and Sholl analysis were exported to separate Excel files and used for data analysis.

**Determination of plasma OT levels by radioimmunoassay (RIA)**

Mice were decapitated and trunk blood (0.5-1 mL) was collected in chilled tubes containing 50 µl of 0.3 M EDTA (pH 7.4) and 100 µl ml of aprotinin (0.6TIU/mL of blood). Samples were stored on ice for a maximum of 15 min before centrifugation at 1,600 x g for 15 min (4°C). Plasma was collected, transferred to polypropylene tubes and stored at -80°C until OT RIA. OT levels were determined from 350 µl of plasma using a specific RIA kit (Phoenix Pharmaceutical, Inc, RK-051-01) using $^{125}$I OT (1227 Ci/mmol) as a tracer and a polyclonal antiserum specific for OT with no cross reactivity with vasopressin, Met-enkephalin, somatostatin and vasointestinal peptide. OT was extracted from the acidified plasma with a Sep-Pak C18 cartridge and measured in duplicate according to the protocol of the kit. Experiments were performed in the afternoon.

**In vivo OT microdialysis**

Microdialyses were performed on +/+ and -/- mice in the afternoon. Implantation of microdialysis probes for monitoring OT release within the SON (U-shaped, molecular cutoff 10 kDa; 0.6 mm caudal to bregma, 1.2 mm lateral to midline, 5.5 mm below the surface of the skull) was performed under isoflurane anesthesia (Forene, Abbott GmbH, Wiesbaden, Germany) according to[7]. To avoid post-surgical infections, mice received subcutaneously administered antibiotics (3 mg/30 ml Baytril, Bayer GmbH, Leverkusen, Germany). After surgery, mice were repeatedly handled before experiments started. Two days after surgery, the microdialysis probe was perfused for 2 h with sterile Ringer solution (pH 7.4; 3.3 ml/min) to establish equilibrium between the inside and outside of the microdialysis membrane. Then, three consecutive dialysates were collected: before (basal conditions), during and after (baseline conditions) pup exposure (1 unfamiliar pup, postnatal days 1-4, presented in the home cage) of pup-naïve female mice. Dialysates were immediately frozen and stored at -20°C until quantification of OT by a radioimmunoassay (sensitivity 0.3 pg / sample; RIAgnosis, Sinzing; Germany)[8]. Correct placement of microdialysis probes was validated by post-hoc histological assessment.

**Electron Microscopy**

***Fixation and tissue processing***. +/+ and -/- mice were used (3 mice per condition). Mice were deeply anaesthetized with ketamin (60 mg/kg body weight) and then transcardially perfused through the ascending aorta using a rotation pump (at a constant flow rate) with 0.1M phosphate buffered (PB) saline for 1-2 min, followed by the ice-cold PB-buffered fixative containing 4% paraformaldehyde and 0.1 or 0.5% glutaraldehyde for 10 min. Afterwards, brains were removed from the skull, post-fixed for 1h in the same but fresh fixative at 4°C, then extensively washed in PB and stored in the same buffer until use.

Coronal (frontal) sections through the SON were cut at 150 µm thickness using a vibratome (VT1000S; Leica Microsystems GmbH, Wetzlar, Germany) and collected in PB. After incubation in sucrose-PB containing 1% osmium tetroxide for 1h, sections were washed in PB, and dehydrated in an ascending series of ethanol (10%, 20%, 30%, 50%, 60%, 70%, 80%, 90%, 96%; 15 min for each step) to absolute ethanol (2x 30 min each). Afterwards, sections were transferred to propylene oxide (twice 2 min each), then to a mixture of propylene oxide and epoxy resin (2:1; 1:1 for 1hr each; Durcupan; ACM, Fluka, Neu-Ulm, Germany) and finally transferred to and stored overnight in pure resin. The next day sections were flat-embedded in fresh Durcupan between coated glass slides, coversliped and polymerized at 60$^{o}$C for 2 days.

From the embedded material, blocks containing the SON were selected after light microscopic inspection and pictures were taken for documentation of the area of interest. Prior to serial ultrathin sectioning, semi-thin sections were cut, toluidine-blue stained and examined with light microscopy to identify the region of interest, the dendritic layer within the SON. Then serial ultrathin sections (~20-30 sections/series; ~55nm in thickness, silver to light gray interference contrast appearance) were cut through the areas of interest with an ultramicrotome (Ultracut S; Leica Microsystems GmbH, Wetzlar, Germany). Sections were collected on Pioloform-coated slot copper grids (Fa. Plano, Wiesbaden, Germany). After counterstaining with 5% uranyl acetate in double distilled water (5-20 min) and lead citrate (2-7 min), grids were examined with a Libra 120 electron microscope equipped with a bottom-mounted ProScan 2K digital camera and the SIS Analysis software (Olympus Soft Imaging Solutions GmbH, Hamburg, Germany). Digital images using the Multi Images Acquisition (MIA) software were taken at various magnifications.

***Quantitative analysis of the glial coverage*.** To quantify the astrocytic coverage in +/+ and -/- mice, the following experimental approach was used. Frames of 5 by 5 images were taken using the MIA function of the SIS software within the dendritic region of the SON in 20-30 consecutive ultrathin sections at a final EM magnification of x 6300. Thereafter, using the interactive software ImageJ, the first, the middle and the last MIA image of a series (n = 3 measurements per animal) in the +/+ (n = 3) and -/- (n = 3) mice were used for a further quantitative volumetric analysis. In each section of a group, a grid (grid size 1 $\mu m^2$) was placed over the MIA and in each square the presence or the lack of fine astrocytic processes was measured throughout the images. Using the Cavalieri method, the absolute (volume) contribution of astrocytic processes was determined according to: Cavalieri Estimator:

$$V = a(p) \times \Sigma P \times t$$

where a(p) is the size of one square (1 $\mu m^2$); P: the number of squares counted; t: the thickness of the slice.

In addition, to estimate the synaptic coverage, the SIS software was used to determine the percentage of astrocytic profiles to the total perimeter of either synaptic boutons or dendrites forming a synaptic complex. Out of a series of 4 images, measurement of 5 dendrites and 5 boutons each (= 20 measurements each per animal) in +/+ (n = 3) and -/- (n = 3) mice were taken for analysis. Furthermore, using the ImageSP software (Fa. Tröndle, Moorenweis, Germany) the distance of astrocytic processes to their nearest active zone was measured (20 measurements each per animal) in +/+ (n = 3) and -/- (n = 3) mice.

***3D volume reconstructions***. To illustrate the astrocytic coverage of synaptic complexes in the dendritic region of the SON, 3D-volume reconstructions were made from the z-stacks for each group. Within the z-stack, synaptic complexes and fine astrocytic processes were outlined in different colors (magenta: fine astrocytic processes, yellow: synaptic boutons, blue: postsynaptic target structures) using the contour mode of OpenCAR[9]. All illustrations were performed offline using a batch version of OpenCAR, which generates the 3D reconstructions.

**Behavioral tests**

***Pup exposure test***. Virgin pup-naïve adult females were removed from their home-cage and isolated in a cage measuring 35 X 19 X 14 cm containing clean bedding. After a 10-minute habituation period, an unfamiliar 2- to 4-day-old +/+ pup was introduced in the middle of the cage for 20 minutes. To control for effects of handling, mice were placed in a clean cage and allowed to habituate for 10 minutes, after which a gloved hand touched the bedding in the middle of the cage to mimic placement of a pup. Subjects then remained in the cage for an additional 20 minutes.

***Maternal care test***. Maternal behaviors were evaluated using adult pup-naïve female mice[10]. Mice were placed in a non-enriched cage measuring 35 X 19 X 14 cm and containing clean sawdust. Before starting the assay, animals were acclimated for 10 min in this new environment. One +/+ pup from postnatal days 2 to 4 was then added to the cage and female-pup social interactions were recorded for 20 minutes. The duration and number of female-pup social interactions were scored from videos using Noldus Observer XT14 software (Noldus Information Technology, Leesburg, VA). The observers were uninformed of the genotypes during scoring. Investigative behaviors of female-pup social interaction included sniffing/licking, nursing or moving the pup. The data was extracted and maternal care was analyzed minute by minute for each group.

For experiments with OTR antagonist, 20 minutes prior to the test, we proceeded to an OTR antagonist IP injection or stereotaxic injection in the MPOA (L368 899 or d(CH2)51,Tyr(Me)2,Thr4,Orn8,des-Gly-NH29)-Vasotocin Trifluoroacetate, respectively). For control, we performed similar injections with a saline solution.

***Nest building test***. For assessing nest building, we used the same protocol that has been previously described[11]. Briefly, pup-naïve female mice were individually housed in a non-enriched cage measuring 33 X 14.5 X 13 cm and containing clean sawdust. Two cotton squares (nesting material) were introduced 2h before the onset of the dark period. The animals were monitored for nest building at different time points (1, 2 and 24h after addition of the nesting material) and scored on a 0-5 scale, with 0 representing no contact with the cotton and 5 a perfectly made nest.

***Pup retrieval test***. For measuring pup retrieval, experiments were adapted from procedures that have been previously described[10]. In short, pup-naïve female mice were placed in a non-enriched cage measuring 35 X 19 X 14 cm and containing clean sawdust as well as nesting material coming from the dam's cage. Animals were given at least 20 min to acclimate before each testing session began. Three to four pups of the same litter from postnatal days 2 to 4 were grouped in a corner of the cage and covered with nesting material. At each trial, one pup was removed from the nest and placed in an opposite corner of the cage. The tested female was given 2 min to retrieve the displaced pup and return it back to the nest. The testing sessions consisted of a set of 10 trials. Each trial was scored as follows: if the displaced pup was retrieved within the 2 min, the trial was scored as a success; otherwise, it was scored as a failure. The percentage of pup retrieval for each testing session was calculated as follows: for a tested mouse that has succeeded to retrieve pups in 5 out of the 10 trials, the percentage of pup retrieval is 50%. At each trial, the displaced pup was returned to the nest and another pup was taken out of the nest and placed in an opposite corner of the cage. After the testing session, pups were placed back into their home cage with their dam. Testing sessions were performed on three consecutive days, 3-4 hours after the onset of the light period.

***Locomotor activity.*** We used 40X 40 X 40 cm white opaque PVC openfield (Noldus, The Netherlands). Each mouse was removed from its home cage and put into a holding box next to the testing box for 10 minutes. Subsequently, the mouse was put into the testing box facing the rear wall. Path tracking has been recorded with Ethovision XT 17 (Noldus, The Netherlands). Locomotor activity was measured for 5 minutes. The time in the center, which was determined as 30 cm away from each wall of the box, was measured automatically, when the center-point of the mouse moved into it.

### Statistics

For statistical comparison, normality test as well as variance analysis were performed with a Shapiro-Wilk test and F-test, respectively, and the appropriate two-sided statistical parametric or non-parametric test was used. Two-tailed unpaired or paired t tests were used for between-group comparisons. Statistical significance for within-group comparisons was determined by one-way or two-way ANOVAs followed by post hoc tests. Appropriate sample sizes were based on best practices in the literature as well as on ethical standards to minimize numbers of animals for experiments, and were dictated by the magnitude of experiment-to experiment variation. Animals were not randomized, and blinding to group assignment was not performed. There was no data exclusion. All statistical analysis was performed in GraphPad Prism (GraphPad Software, USA).

**Supplementary figures and legends**

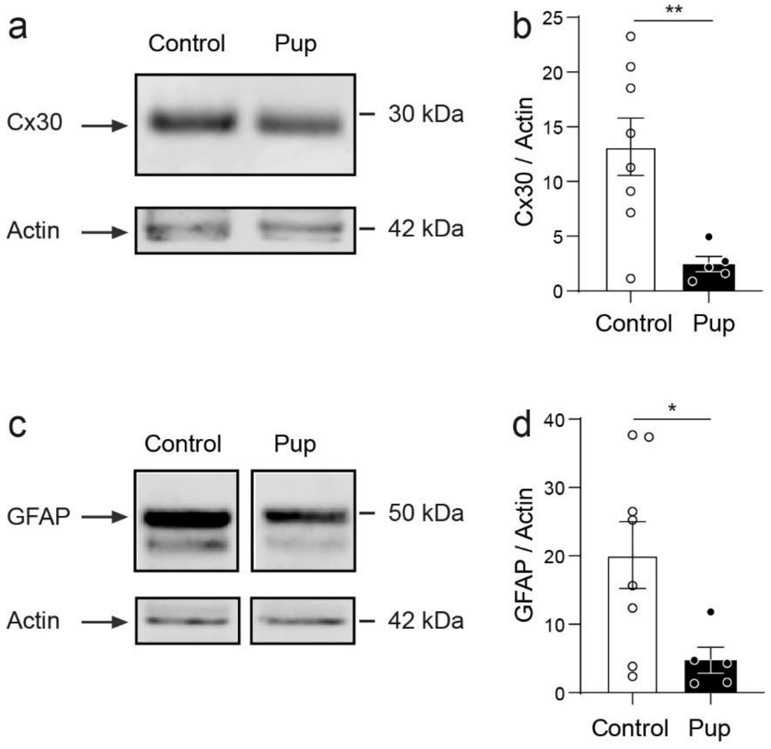


**Supplementary Fig. 1. Pup exposure decreases Cx30 and GFAP levels in the SON of virgin pup naïve female. a.** Western blots showing Cx30 protein expression in the SON of virgin pup-naïve and pup-sensitized female +/+ mice. Actin was used as a loading control. **b.** Quantification showing a decrease in Cx30 levels in the SON of female +/+ mice following pup exposure (control n = 8 samples, pup n = 5 samples, p = 0.0095, unpaired t test). **c.** Western blot showing GFAP protein expression in the SON of virgin pup-naïve and pup-sensitized female +/+ mice. Actin was used as a loading control. **b.** Quantification showing a decrease in GFAP levels in the SON of female +/+ mice following pup exposure (control n = 8 samples, pup n = 5 samples, p = 0.0360 unpaired t test). 8 SON were pooled for each sample (bilateral punches from 4 mice). Asterisks indicate statistical significance (*p < 0.05; **p < 0.01).

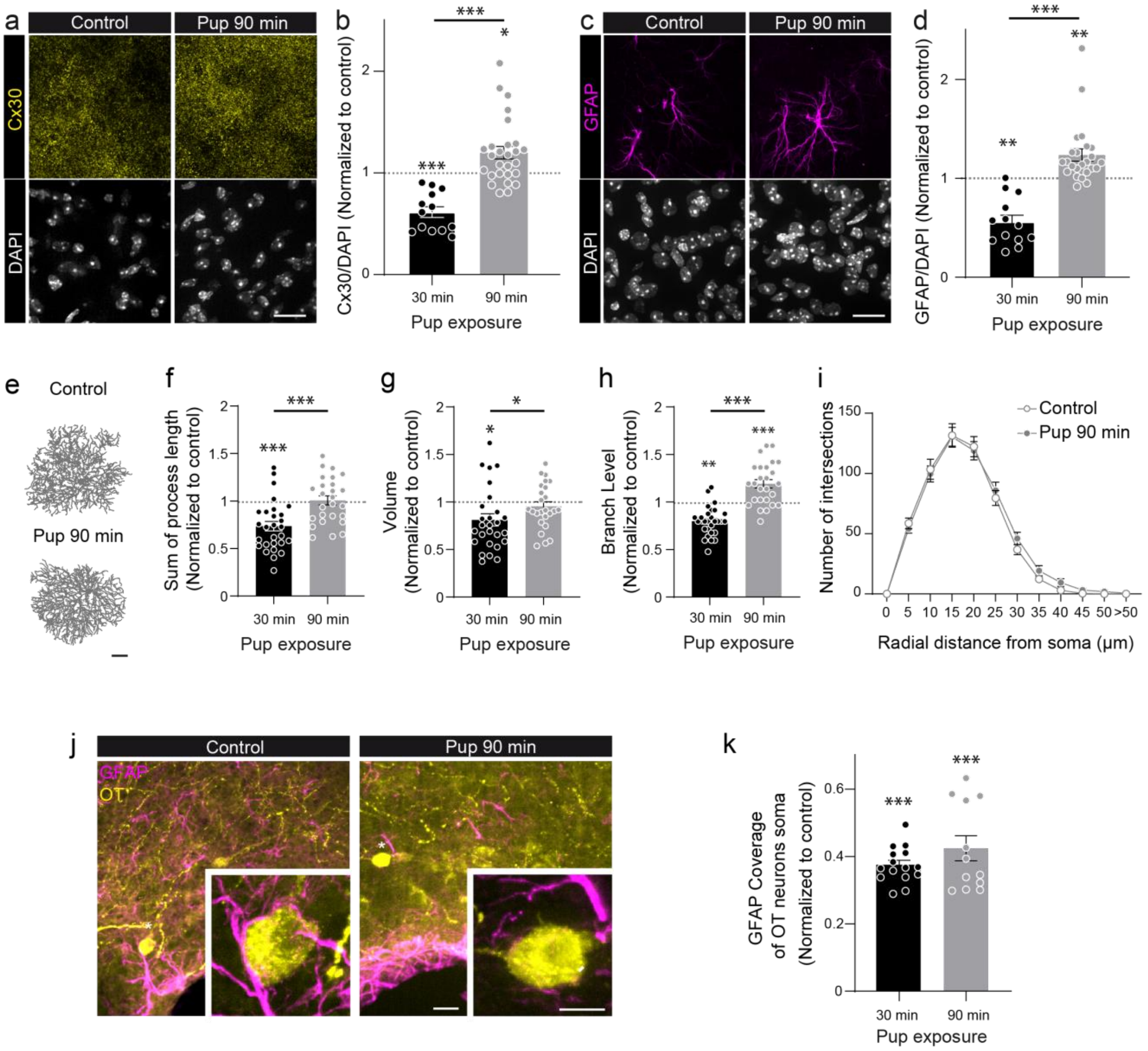


**Supplementary Fig. 2. Astrocyte morphological and synapse coverage changes mediated by pup exposure are transient. a** Representative images showing Cx30 (yellow) and DAPI (white) fluorescent immunostainings in the SON of virgin pup-naïve vs pup-sensitized female +/+ mice sacrificed 90 minutes after the beginning of the test. Scale bar: 30 µm. **b** Pup-sensitized female +/+ mice sacrificed 90 minutes after the beginning of the test (grey, n=27 ROI from 6 mice) show an increase in Cx30 fluorescence intensity compared to pup-sensitized female +/+ mice sacrificed after 30 minutes (black, n = 13 ROI from 5 mice, $p < 0.0001$, Mann Whitney test) Fluorescence intensity of Cx30 was normalized to DAPI fluorescence intensity and the values from pup-sensitized mice were normalized to values from pup-naïve mice. **c** Representative images showing GFAP (magenta) and DAPI (white) fluorescent immunostainings in the SON of virgin pup-naïve vs pup-sensitized female +/+ mice sacrificed 90 minutes after the beginning of the test. Scale bar: 30 µm. **d** Pup-sensitized female +/+ mice sacrificed 90 minutes after the beginning of the test (grey, n=24 ROI from 6 mice) show an increase in GFAP fluorescence intensity compared to pup-sensitized female +/+ mice sacrificed after 30 minutes (black, n = 13 astrocytes from 5 mice, $p < 0.0001$, Mann Whitney test). Fluorescence intensity of GFAP was normalized to DAPI fluorescence intensity and the values from pup-sensitized mice were normalized to values from pup-naïve mice. **e** Three-dimensional reconstruction of astrocytes in the SON of virgin pup-naïve (up) vs pup-sensitized (bottom) female +/+ mice sacrificed 90 minutes after the beginning of the test. **f** Restoration of the sum of process length of astrocytes in the SON of pup-sensitized female +/+ mice sacrificed 90 minutes after the beginning of the test (grey, n= 26 astrocytes from 6 mice) vs pup-sensitized female +/+ mice sacrificed 30 minutes after the beginning of the test (black, n= 28 astrocytes from 4 mice, $p = 0.0003$, unpaired t-test). **g** Restoration of domain volume of astrocytes in the SON of pup-sensitized female +/+ mice sacrificed 90 minutes after the beginning of the test (grey, n= 25 astrocytes from 6 mice) vs pup-sensitized female +/+ mice sacrificed 30 minutes after the beginning of the test (black, n= 27 astrocytes from 4 mice, $p = 0.0246$, Mann Whitney test). **h** Increase of the branch level of astrocytes in the SON of pup-sensitized female +/+ mice sacrificed 90 minutes after the beginning of the test (grey, n= 26 astrocytes from 6 mice) vs pup-sensitized female +/+ mice sacrificed 30 minutes after the beginning of the test (black, n= 28 astrocytes from 4 mice, $p < 0.0001$, unpaired t-test). **i** Restoration of the number of intersections per radial distance from the soma determined by Sholl analysis of astrocytes in the of pup-sensitized (grey, n= 26 astrocytes from 6 mice) compared to pup-naïve female +/+ mice sacrificed 90 minutes after the beginning of the test vs pup-sensitized female +/+ mice sacrificed 30 minutes after the beginning of the test (black, n= 24 astrocytes from 4 mice, $p=0.3767$, two-way ANOVA). **j** Representative images of oxytocinergic neurons (yellow) covered by GFAP-rich astrocyte processes (magenta) in the SON of virgin pup-naïve vs pup-sensitized female +/+ mice sacrificed 90 minutes after the beginning of the test. The white asterisks indicate neurons that are magnified in the insets. Scale bar: 20 µm; inset: 8 µm. **k** The coverage of oxytocinergic neurons remains reduced in pup-sensitized female +/+ mice sacrificed 90 minutes after the beginning of the test (grey, n = 12 astrocytes from 3 mice) and was similar to the one of pup-sensitized female +/+ mice sacrificed 30 minutes after the beginning of the test (black, n = 15 astrocytes from 3 mice, $p = 0.1925$, unpaired t-test). Asterisks indicate statistical significance (*$p < 0.05$; **$p < 0.01$; ***$p < 0.001$).

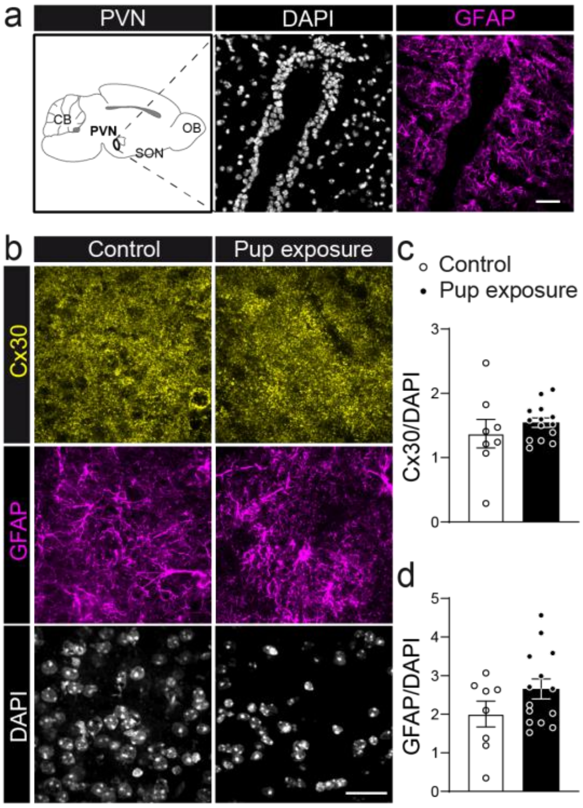


**Supplementary Fig. 3. Pup exposure does not alter Cx30 expression in the PVN of virgin pup-naïve female mice. a** Scheme depicting localization of the SON and representative images of DAPI (cyan) and GFAP (magenta) double-immunofluorescence labeling in the PVN of +/+ mice. Scale bar = 30 µm. **b** Representative images showing Cx30 (yellow), GFAP (magenta) and DAPI (white) fluorescent immunostainings in the PVN of virgin pup-naïve vs pup-sensitized female +/+ mice. Scale bar = 25 µm **c, d** Female +/+ mice sensitized to pups displayed unchanged Cx30 (n = 14 ROI from 6 mice, p= 0.38, unpaired t-test) and GFAP (n = 14 ROI from 6 mice, p = 0.1414, unpaired t-test) fluorescence intensities compared to pup-naïve controls (n = 8 ROI from 3 mice). Fluorescence intensities of Cx30 and GFAP are normalized to DAPI fluorescence intensity.

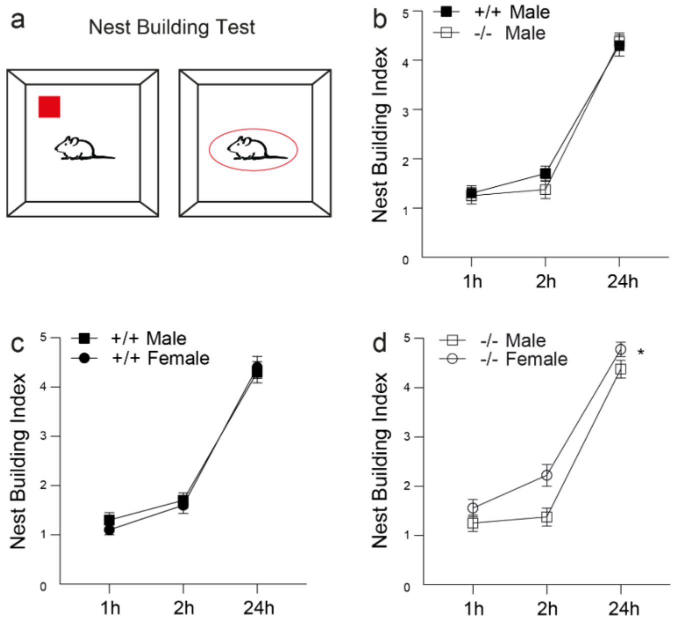


**Supplementary Fig. 4. Sexually dimorphic control of nest building behavior by Cx30. a** Illustration of the nest building test used to assess parental care in mice. **b** Pup naïve -/- male mice (n = 8) show similar nest building performances as compared to pup naïve +/+ males (n = 10, p = 0.627, two-way repeated measures ANOVA). **c** In pup naïve +/+ mice, both males (n = 10) and females (n = 10) showed similar nest-building capacities (p = 0.708, two-way repeated measures ANOVA). **d** In pup naïve -/- mice, males (n = 8) display qualitatively lower nest building activity compared to females (n = 9, p = 0.0147, two-way repeated measures ANOVA). Asterisks indicate statistical significance (*p < 0.05).

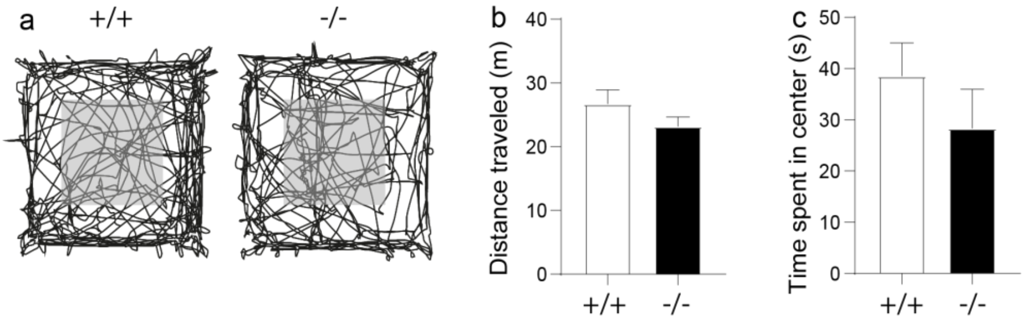


**Supplementary Fig. 5. Astroglial Cx30 deficiency has no effect on locomotion and anxiety.** **a** Representative travel paths of +/+ and -/- female pup-naïve mice performing an open-field test. Grey squares represent the center of the arena. **b** The distance traveled by -/- virgin pup-naïve female mice (n=8) is similar to +/+ female pup-naïve mice (n=6, p= 0.1928 unpaired t-test). **c** The time spent in the center by -/- virgin pup-naïve female mice (n=8) is similar to +/+ virgin pup-naïve female mice (n=6, p=0.3491 unpaired t-test).

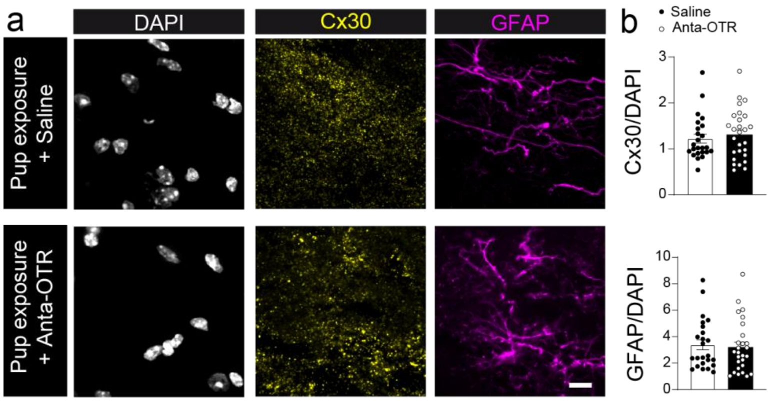


**Supplementary Fig. 6. OTR antagonist does not affect changes in the expression of Cx30 and GFAP triggered by pup exposure in the SON of virgin pup-naïve female mice. a** Representative images showing Cx30 (yellow), GFAP (magenta) and DAPI (white) fluorescent immunostainings in the SON of pup-sensitized female +/+ mice injected with anta-OTR or saline solution. Scale bar: 10 µm. **b** Female +/+ mice sensitized to pups injected with anta-OTR displayed unchanged Cx30 (n = 22 ROI from 5 mice, p =8955, unpaired t-test) and GFAP (n = 22 ROI from 5 mice, p =5554, Mann Whitney test) fluorescence intensities compared to pup-sensitized female +/+ mice injected with saline (n = 21 ROI from 5 mice). Fluorescence intensities of Cx30 and GFAP are normalized to DAPI fluorescence intensity.

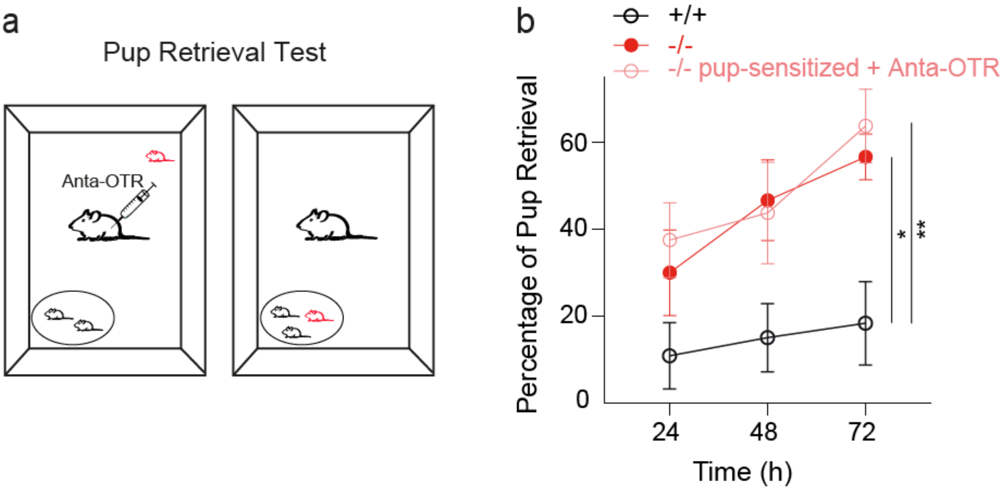


**Supplementary Fig. 7. Pup retrieval after maternal learning is independent of OT signaling in -/- mice. a** Schematic representation of the pup retrieval test used to assess pup-directed maternal care in pup-naïve vs pup-sensitized -/- mice injected intra-peritoneally with an OTR antagonist (Anta-OTR, L368 899). **b** Pup-sensitized -/- female mice injected with the OTR antagonist (n = 8) show similar pup retrieval capacities compared to virgin pup naïve -/- female mice (n = 9, p = 0.702, two-way ANOVA repeated measures) and was higher than capacities of virgin pup naïve +/+ female mice (n = 10, p=0.0088 and p=0.0150, respectively, two-way ANOVA repeated measures). Asterisks indicate statistical significance (*p < 0.05).

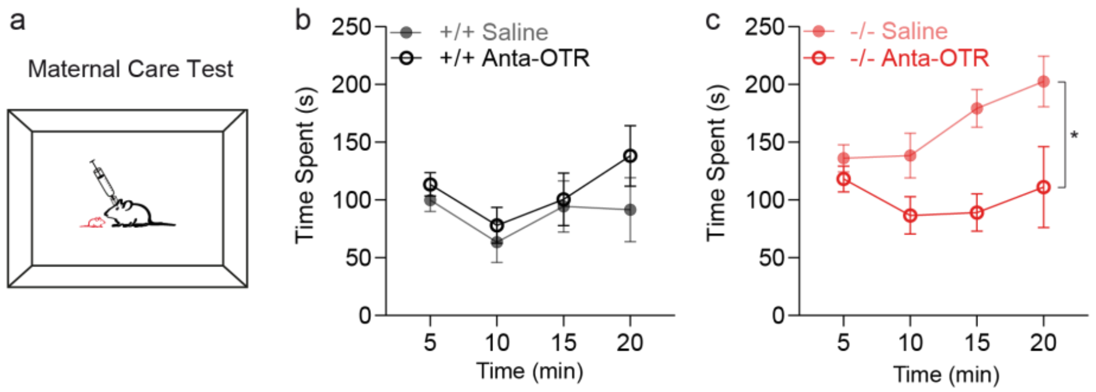


**Supplementary Fig. 8. Cx30 controls maternal behavior through OT signaling in the MPOA. a** Schematics showing the experimental protocol applied to measure the time spent in active maternal care in virgin pup naïve +/+ and -/- female mice stereotaxically injected via a cannula in the MPOA with saline or an OTR antagonist (Anta-OTR, d(CH2)51,Tyr(Me)2,Thr4,Orn8,Tyr-NH29)-vasotocin trifluoroacetate salt). **b** The time spent with pups in virgin pup naïve female +/+ mice injected in the MPOA with the OT antagonist (n = 8) did not differ from saline control virgin pup naïve female +/+ mice (n = 8, p = 0.3913, two-way repeated measures ANOVA). **c** In contrast, virgin pup naïve female-/- mice injected in the MPOA with the OTR antagonist (n = 7) differs significantly from saline control -/- mice (n = 6, p = 0.0155, two-way repeated measures ANOVA), indicating that Cx30 sets maternal care through OT signaling in the MPOA. Asterisks indicate statistical significance (*p < 0.05; **p < 0.01; ***p < 0.001).

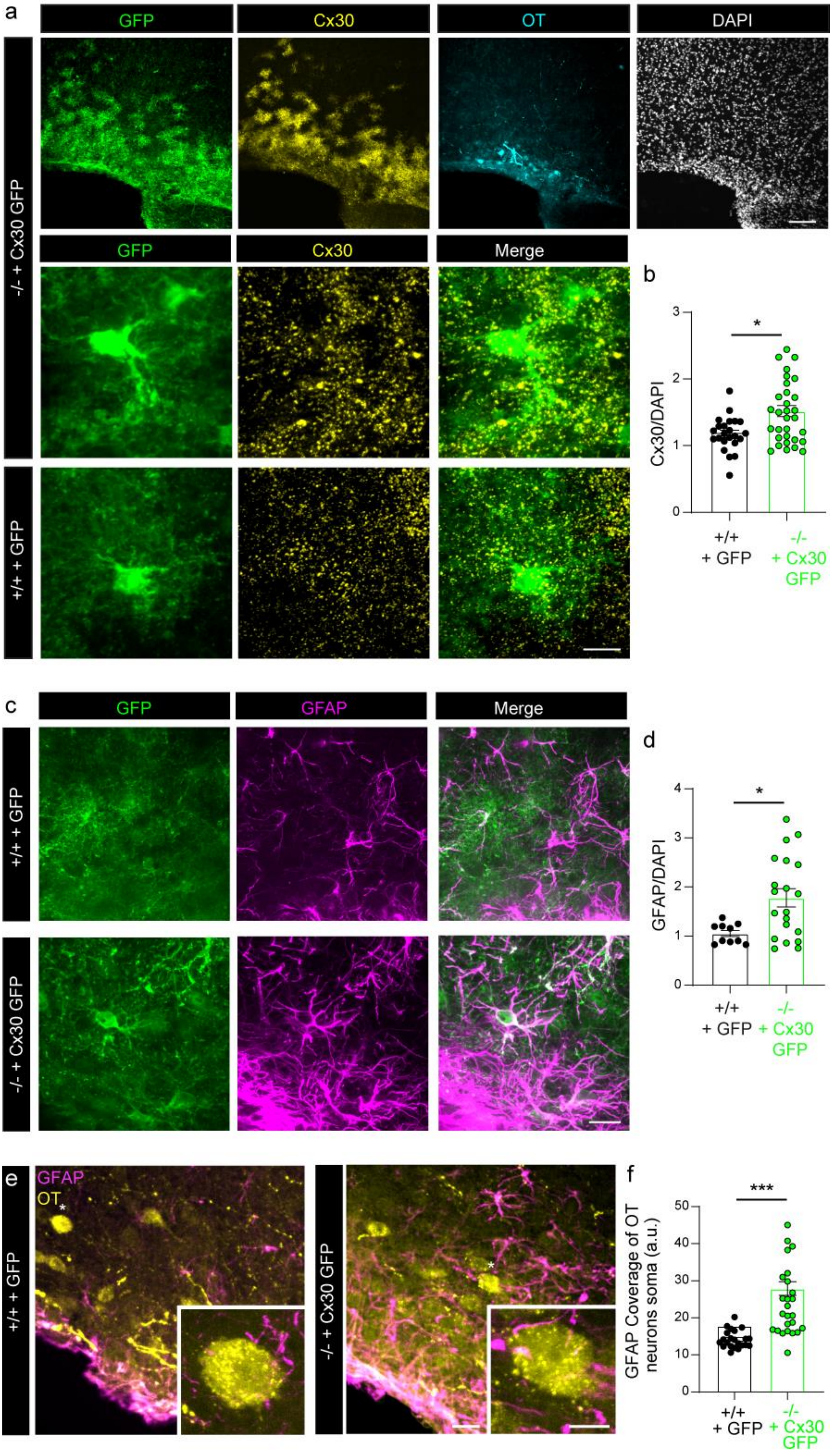
a
GFP
Cx30
OT
DAPI
-/- + Cx30 GFP
GFP
Cx30
Merge
+/+ + GFP
b
Cx30/DAPI
*
+/+
+ GFP
-/-
+ Cx30
GFP
c
GFP
GFAP
Merge
+/+ + GFP
-/- + Cx30 GFP
d
GFAP/DAPI
*
+/+
+ GFP
-/-
+ Cx30
GFP
e
GFAP
OT
+/+ + GFP
-/- + Cx30 GFP
f
GFAP Coverage of OT neurons soma (a.u.)
***
+/+
+ GFP
-/-
+ Cx30
GFP

**Supplementary Fig. 9. Restoring Cx30 locally in astrocytes increases astroglial coverage of OT neurons in the SON of virgin female -/- mice. a** Representative images showing the restoration of Cx30 expression (yellow) in the SON of virgin pup naïve female -/- mice at low and high magnification (scale bar: 50 and 10 µm). GFP (green), Cx30 (yellow), oxytocinergic neurons (cyan) and DAPI (white) immunolabelling in the SON of virgin pup naïve female -/- mice locally injected with AAV encoding Cx30 and GFP and virgin pup naïve +/+ mice locally injected with AAV encoding GFP. **b** The expression of Cx30 is increased in the SON of virgin pup naïve female -/- mice locally injected with AAV encoding Cx30 and GFP (n=31 ROI from 5 mice) compared to +/+ mice locally injected with AAV encoding GFP (n=23 ROI from 8 mice, p=0.0210 Mann Whitney test). Fluorescence intensity of Cx30 is normalized to DAPI fluorescence intensity. **c** Representative images showing GFP (green), and GFAP (magenta) fluorescent immunostainings in the SON of virgin pup naïve female -/- mice locally injected with AAV encoding Cx30 and GFP, and to +/+ mice locally injected with AAV encoding GFP. Scale bar: 20 µm. **d** The expression of GFAP is increased in the SON of virgin pup naïve female -/- mice locally injected with AAV encoding Cx30 and GFP (n=20 slices from 6 mice) compared to +/+ mice locally injected with AAV encoding GFP (n=10 slices from 6 mice, p=0.0113 unpaired t-test). Fluorescence intensity of GFAP is normalized to DAPI fluorescence intensity. **e** Representative images of oxytocinergic neurons (yellow) covered by GFAP-rich astrocyte processes (magenta) in the SON of virgin pup naïve female -/- mice locally injected with AAV encoding Cx30 and GFP, and to +/+ mice locally injected with AAV encoding GFP. The white asterisks indicate neurons that are magnified in the insets. Scale bar: 20 µm; inset: 8 µm. **f** The coverage of oxytocinergic neurons is increased in virgin pup naïve female -/- mice locally injected with AAV encoding Cx30 and GFP (green, n = 25 astrocytes from 6 mice) compared to +/+ mice locally injected with AAV encoding GFP (black, n = 22 astrocytes from 6 mice, $p < 0.0001$, unpaired t-test). Asterisks indicate statistical significance (*$p < 0.05$; ***$p < 0.001$).

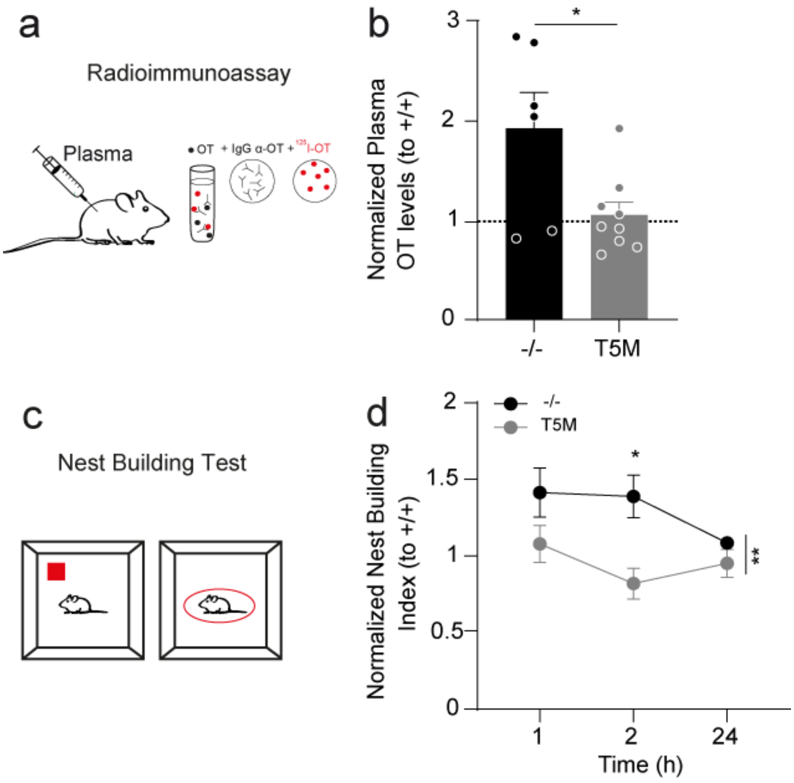


**Supplementary Fig. 10. Astroglial Cx30 in the SON regulates plasma OT levels and maternal behavior independently of gap junction-mediated biochemical coupling. a** Schematic representation of the radioimmunoassay (RIA) method used to measure OT plasma levels in mice. **b** Plasma OT levels measured by RIA were decreased in virgin pup naïve female T5M mice (grey, n = 9) compared to -/- mice (black, n = 6, p = 0.0211, unpaired t-test) and was similar to +/+ mice (n = 8, p = 0.7118, unpaired t-test. **c** Schematic representation showing nest building procedure used to assess maternal care in virgin pup naïve female T5M mice. **d** Normalized performances of virgin pup naïve female T5M mice (n = 8) differs significantly from that of -/- mice (n = 9, p = 0.0069, two-way repeated measures ANOVA), thus indicating that Cx30 controls maternal behavior and OT signaling independently of gap junction-mediated biochemical coupling. Asterisks indicate statistical significance (*p = 0.05; **p < 0.01).
